# The Nuclease Domain of *E. coli* RecBCD Regulates DNA Binding and Base Pair Melting

**DOI:** 10.1101/2025.07.11.664400

**Authors:** Linxuan Hao, Rui Zhang, Timothy M. Lohman

## Abstract

*E. coli* RecBCD, a hetero-trimeric helicase and nuclease, functions in double stranded (ds) DNA break repair and in degrading foreign DNA. RecBCD possesses ATPase motor domains within both RecB (3’ to 5’) and RecD (5’ to 3’) and a nuclease domain within RecB (RecB^Nuc^). RecBCD binds to double stranded DNA ends and initiates DNA unwinding by first melting several DNA base pairs (bp) using only its binding free energy. The RecB^Nuc^ domain is docked ∼70 Å from the duplex DNA binding site in RecBCD-DNA structures but appears to be dynamic and able to move from its docked position. Here, we compare DNA binding of RecBCD and a variant, RecB^ΔNuc^CD, in which the 30 kDa nuclease domain has been deleted. RecBCD binding to a blunt DNA end is enthalpically unfavorable and entropically driven. Deletion of RecB^Nuc^ results in an increase in DNA binding affinity, suggesting an allosteric effect of RecB^Nuc^. RecB^ΔNuc^CD binding to DNA possessing fully ‘pre-melted’ DNA ends is associated with a large favorable ΔH_obs_, but much smaller than observed for RecBCD, suggesting that deletion of RecB^Nuc^ limits bp melting from a blunt DNA. We also solved cryo-EM structures showing only 4 bp melted upon RecB^ΔNuc^CD binding to a blunt ended DNA duplex, less than the 11 bp melted upon RecBCD binding. Thus, the RecB nuclease domain regulates the extent of bp melting by RecBCD. These results also suggest that RecB^Nuc^ may manifest its long-range allosteric effect on DNA binding and DNA melting via linker-linker interactions between RecB and RecC.

## Introduction

Double strand (ds) DNA breaks in the chromosome are lethal if not repaired. Repair of dsDNA breaks in *E. coli* relies primarily on homologous recombinational repair ^1–4^. Recombinational repair in *E. coli* is initiated when the RecBCD helicase/nuclease binds to the broken dsDNA end and unwinds the dsDNA using its helicase activity. *E. coli* RecBCD is a complex molecular machine composed of three subunits. RecB contains a canonical superfamily (SF) 1A (3’ to 5’) translocase motor that is tethered via a ∼60 amino acid linker to a C-terminal nuclease domain (RecB^Nuc^), RecD contains a canonical SF1B (5’ to 3’) translocase motor^5–8^ and RecC is a processivity factor that interacts with both RecB and RecD and contains the site for recognition of the crossover hotspot initiator (chi) site (5’-GCTGGTGG)^9,10^, which serves to identify the DNA as host chromosomal DNA chi sequence. A secondary translocase activity, associated with the RecB “arm” domain, functions within either RecBC or RecBCD, and is controlled by the RecB ATPase and is likely a dsDNA translocase activity^11–13^.

Upon initial binding to a blunt dsDNA end, RecBCD uses its binding free energy, in the absence of ATP to destabilize (“melt”) several base pairs (bp), ranging from 4-6 bp^14–16^, to 11 DNA bp^17^ depending on the DNA duplex length. This initiation step produces the ssDNA that interacts directly with the RecB motor domain that then initiates processive DNA unwinding coupled to ATP binding and hydrolysis^18,19^. While carrying out DNA unwinding, the RecB nuclease domain degrades both strands of the unwound single stranded (ss) DNA^20,21^. However, a series of changes occur when RecBCD encounters and recognizes a chi site. These include a ∼two-fold reduction in the DNA unwinding rate, a change in the relative rates of translocation of the RecB and RecD canonical motors, as well as a change in the nuclease activity such that only the 5’-ended ssDNA is degraded^22–27^. RecA protein is subsequently loaded onto the resected 3’-ended ssDNA and this has been proposed to occur via direct interactions of RecA with the nuclease domain^22,28,29^. The RecA coated 3’-ended ssDNA then undergoes a search for sequence homology to initiate DNA repair via DNA recombination.

In all RecBCD-DNA structures in which the nuclease domain is visible, it is located in a docked position at the opposite end of the enzyme from the DNA binding site^7,15,17,30,31^. Furthermore, the region of the nuclease domain proposed to interact with RecA protein, is buried in the RecBCD structures^28^. These observations have led to the suggestion that the nuclease domain is dynamic and can move from its docked position in order to load RecA protein^28^. The dynamic nature of the nuclease domain is supported by cryo-EM structures of RecBCD bound to a blunt dsDNA end that show multiple structural classes^17^. We observed previously that one class of RecBCD-DNA shows the nuclease domain in its docked state, whereas a second class shows no density for the nuclease domain, even though the nuclease domain is present in the protein samples^17^.

Fazio et al.^19^ showed that deletion of the RecB nuclease domain to form RecB^ΔNuc^CD inhibits its ability to initiate DNA unwinding from a blunt DNA end and also reduces its rate of DNA unwinding, but not the rate of ssDNA translocation. Based on this, it was proposed that the nuclease domain may regulate the rate or extent of DNA bp melting. In order to probe the effect of the nuclease domain on RecBCD binding and DNA bp melting, we have compared the thermodynamics of DNA binding of RecBCD with RecB^ΔNuc^CD, complemented by Cryo-EM structures of these enzymes bound to DNA ends. Our results show that the nuclease domain regulates both the energetics of DNA binding and the amount of DNA bp melting upon binding to a blunt DNA end, likely by an allosteric mechanism. This further supports that the nuclease domain plays a regulatory role in the DNA binding and bp melting by RecBCD, consistent with its role in DNA helicase activity.

## Results

### Deletion of the RecB nuclease domain increases binding affinity to DNA ends

RecB^ΔNuc^CD is a variant of RecBCD in which the RecB nuclease domain, RecB^Nuc^ (RecB930-1180) has been deleted. We purified RecB^ΔNuc^CD to homogeneity as described^13^ (see Materials and Methods). A denaturing SDS polyacrylamide gel of purified RecBCD with purified RecB^ΔNuc^CD is shown in Figure 1A. The RecB^ΔNuc^CD sample contains wild type RecC and RecD and RecB^ΔNuc^ runs at the expected lower molecular weight (105 kDa) with no evidence of full length RecB (134 kDa). Unless otherwise specified, experiments were performed in Buffer Mx-y, which is 20 mM MOPS, pH7.0, where ‘x’ indicates the concentration of NaCl in mM and ‘y’ indicates the concentration of MgCl_2_ in mM.

**Figure 1.**
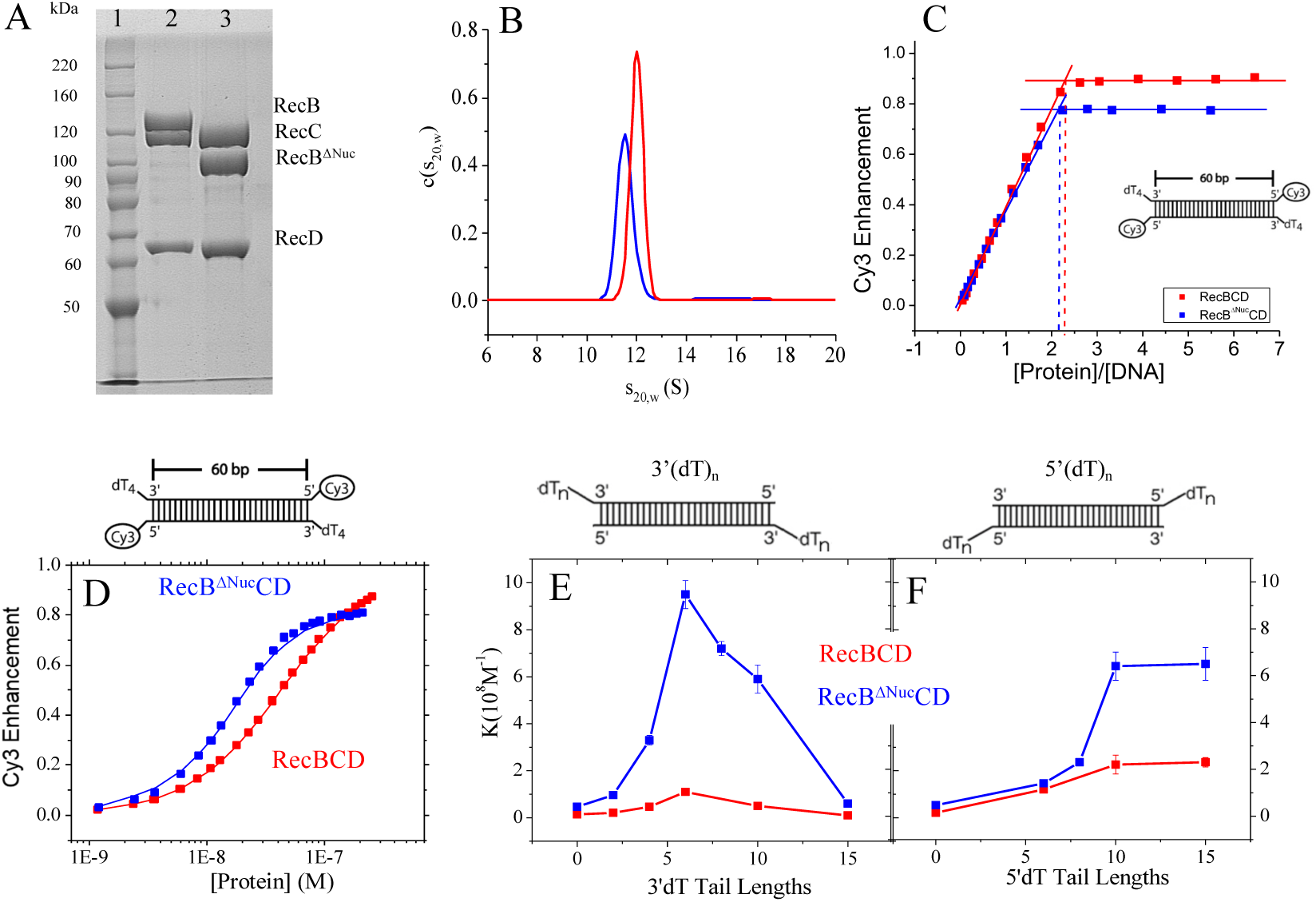
RecB^ΔNuc^CD binds DNA ends with higher affinity than RecBCD. (A)-SDS polyacrylamide (8%) gel of urified RecBCD and RecB^ΔNuc^CD proteins: Lane 1-Benchmark^TM^ protein ladder; Lane 2-RecBCD; Lane 3-RecB^ΔNuc^CD. (B)-Continuous sedimentation, c(s), distribution of 200 nM of RecBCD (red, s_20,w_=11.9 S) and RecB^ΔNuc^CD (blue, s_20,w_=11.5 S) obtained from a sedimentation velocity experiment (Buffer M50-10, 25°C). (C)-Titrations of Cy3 labeled reference DNA with RecB^ΔNuc^CD (blue) and RecBCD (red) (Buffer M50-10, 25°C), showing a stoichiometry of two RecB^ΔNuc^CD or two RecBCD hetero-trimers per DNA molecule. (D)-Fluorescence titration of Cy3 labeled reference DNA with RecB^ΔNuc^CD (blue) and RecBCD (red) (Buffer M275-10, 25°C). (E)-Equilibrium constants for RecBCD (red) and RecB^ΔNuc^CD (blue) binding to unlabeled DNA ends as a function of length (n) of 3’-dT_n_. (F)-Equilibrium constants for RecBCD (red) and RecB^ΔNuc^CD (blue) binding to unlabeled DNA ends as a function of length (n) of 5’-dT_n_.

Over the concentration ranges and in the solution conditions used in this study, RecBCD and RecB^ΔNuc^CD exist as stable hetero-trimers. Figure 1B shows the c(s) distribution from sedimentation velocity experiments in Buffer M50-10, 25°C. Both RecB^ΔNuc^CD and RecBCD samples show single symmetric peaks with s_20,w_= 11.5 S and s_20,w_=11.9 S, respectively. Our RecBCD samples contain only the trimeric form of RecBCD, after removal of the hexameric form of RecBCD^32^. In contrast, we have never detected any hexameric form of RecB^ΔNuc^CD at any point in the purification. This suggests that the nuclease domain is responsible for formation of the (RecBCD)_2_ hexamer and we hypothesize that it is due to a domain swap of the nuclease domain between two hetero-trimers.

We first examined binding of RecB^ΔNuc^CD to DNA ends using a fluorescently labeled reference DNA as described^16,33,34^. This reference DNA consists of a 60 bp duplex with a 5’-Cy3 fluorescent label and a 3’-dT_4_ tail at each end (Supplementary Table 1). The Cy3 fluorescence is enhanced upon RecB^ΔNuc^CD binding. Previous studies^16,33,34^ have shown that this reference DNA is long enough to allow the binding of one RecBCD hetero-trimer per DNA end. Figure 1C shows a binding isotherm from titration of RecB^ΔNuc^CD with reference DNA in Buffer M50-10 at 25°C. The Cy3 fluorescence increases linearly with RecB^ΔNuc^CD concentration and reaches a plateau at ∼80% enhancement with a sharp breakpoint at [RecB^ΔNuc^CD]/[DNA]=2.1 indicating that two RecB^ΔNuc^CD trimers bind to the reference DNA (one to each end) as expected, the same as we observe for RecBCD (Figure 1C). Therefore, our RecB^ΔNuc^CD protein preparation is 100% active in DNA binding. However, the binding affinity of RecB^ΔNuc^CD to the reference DNA is too high (K_BΔNucCD_ > 2×10^9^M^-1^) to be measured accurately under these solution conditions.

To obtain a more accurate measure of the equilibrium constant for binding of RecBCD and RecB^ΔNuc^CD to the reference DNA we increased the NaCl concentration to 275 mM (Buffer M275-10) 25°C and the results are shown in Figure 1D. Surprisingly, RecB^ΔNuc^CD binds with ∼five-fold higher affinity (K_BΔNucCD_ = 1.6(±0.1)×10^8^M^-1^) than RecBCD (K_BCD_ = 3.2(±0.1)×10^7^M^-1^). Hence, the presence of the nuclease domain decreases the affinity of RecBCD to a DNA end.

We next examined RecBCD and RecB^ΔNuc^CD binding to dsDNA ends as a function of the lengths of 3’ or 5’ ssDNA tails in Buffer M275-10, at 25°C. The DNA substrates used (3’-dT_n_ and 5’-dT_n_ in Supplementary Table 1) are not fluorescently labeled. They have the same sequence in the duplex region as the reference DNA but possessed different lengths of either a 3’dT_n_ or 5’dT_n_ tail. The binding experiments were performed using a competitive binding approach using the fluorescent reference DNA shown in Figure 1D as a competitor as described in previous studies of RecBCD^16^. The resulting equilibrium binding constants are shown in Figure 1E, 1F and Supplementary Table 2. The binding isotherms of these competition binding experiments are shown in Supplementary Figure 1.

The binding constants of RecB^ΔNuc^CD (K_BΔNucCD_) to a DNA end increases as the 3’-dT_n_ tail length increases from 0 to 6 nucleotides, but then decreases for n>6 (Figure 1E). K_BΔNucCD_ increases with increasing 5’-dT_n_ tail length, reaching a plateau at n=10 (Figure 1F). These trends for K_BΔNucCD_ are the same as observed for K_BCD_ in previous studies^34^, although in a different buffer condition.

Importantly, K_BΔNucCD_ are consistently higher than K_BCD_ for both 3’-dT_n_ and 5’-dT_n_ substrates across all dT tail lengths. Hence, the presence of the nuclease domain decreases the affinity of RecBCD for a DNA end. In all RecBCD structures published to date^15,31,35,36^, the RecB nuclease domain, RecB^Nuc^, when observed, is located far from the DNA binding site of RecBCD and shows no direct interaction with the DNA, hence this result suggests that the effect of RecB^Nuc^ on DNA binding is allosteric.

### Dissecting the energetics of RecB^ΔNuc^CD-DNA binding from DNA base pair melting

To understand the energetics of RecB^ΔNuc^CD binding to and melting of dsDNA ends we used the approach summarized in Figure 2A as used in our previous study of RecBCD^17^. The total free energy change for RecB^ΔNuc^CD binding to a blunt DNA end, ΔG_°blunt_, is the sum of the favorable contributions from protein-DNA interactions, ΔG_°PD,_ and the unfavorable contributions melting some number of base pairs of duplex DNA, ΔG_°melt,_ as described in Eq. (1a).

**Figure 2.**
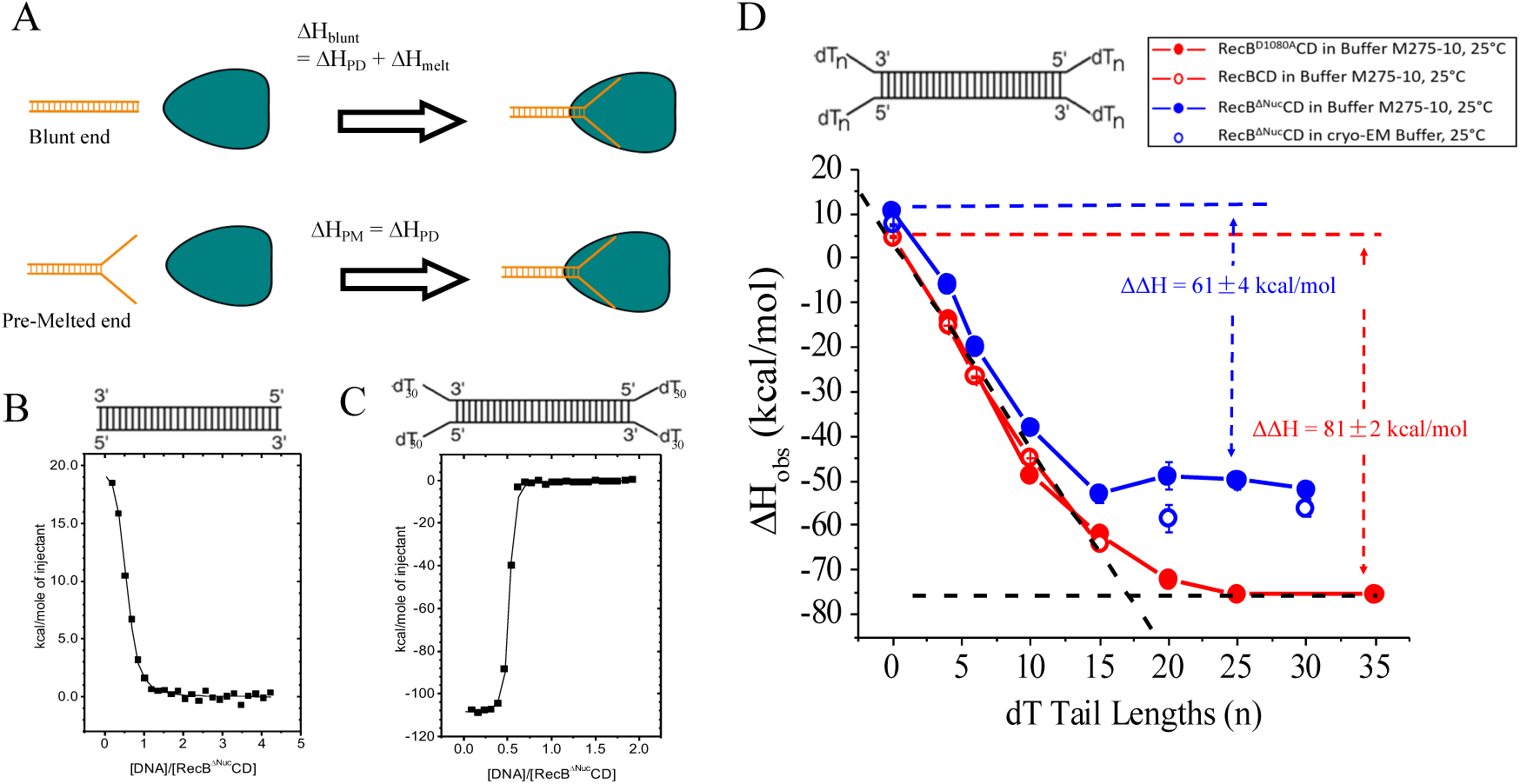
RecB^ΔNuc^CD melts fewer base pairs from a blunt DNA end than RecBCD. (A)-Cartoon depiction of RecBCD or RecB^ΔNuc^CD (green triangle) binding to a blunt DNA end (ΔH_°blunt_ = ΔH_°PD_ + ΔH_°melt_) vs. a fully pre-melted DNA end ((ΔH_°PM_ = ΔH_°PD_). The difference, ΔΔH° = (ΔH_°blunt_ -ΔH_°PM_) = ΔH_°melt_. (B)-ITC experiment for binding of blunt-ended DNA (5 μM) to RecB^ΔNuc^CD (220 µM) in Buffer M275-10, 25°C. (C)- ITC experiment for binding of blunt-ended DNA substrate (5 μM) to RecB^ΔNuc^CD (480 µM) in Buffer M275-10, 25°C. (D)- ΔH_°obs_ for RecBCD (open red circles), RecB^D1080A^CD (filled red circles) and RecB^ΔNuc^CD (filled blue circles) binding to a series of DNA molecules possessing dT_n_-dT_n_ ends of varying lengths (n) (Buffer M275-10, 25°C). RecB^ΔNuc^CD (open blue circles) binding to DNA with blunt, dT_20_- dT_20_ or dT_30_-dT_30_ ends in cryo-EM buffer, 25°C. For RecBCD, the ΔΔH°(BCD) = ΔH_°blunt_ −ΔH_°PM_ (plateau) =81±2 kcal/mol and for RecB^ΔNuc^CD, ΔΔH° (B^ΔNuc^CD) = 61±4 kcal/mol.

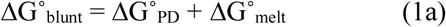

The same relation holds for the enthalpy change associated with DNA binding (Eq. 1b).

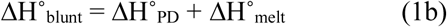

The unfavorable contributions due to base pair melting will be eliminated for RecB^ΔNuc^CD binding to a DNA end that is already fully pre-melted as indicated in Eqs. (2), where the subscript “PM” refers to RecB^ΔNuc^CD binding to a pre-melted DNA end.

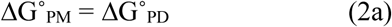

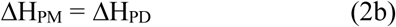

With the assumption that the final state for RecB^ΔNuc^CD bound to a pre-melted DNA end is the same as the final state for RecB^ΔNuc^CD bound to a melted blunt DNA end, we can estimate of the energetic contributions from bp melting by comparing the energetics of RecB^ΔNuc^CD binding to a blunt end vs. a fully melted end as in Eqs. (3).

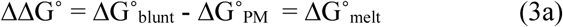

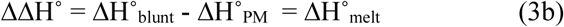

### RecB^ΔNuc^CD melts fewer base pairs than RecBCD upon binding a blunt DNA end

We used isothermal titration calorimetry (ITC) to examine RecB^ΔNuc^CD binding to a 60 bp blunt ended DNA as well as to a 60 bp duplex with DNA ends possessing twin dT_n_ tails of the same length on both the 3’- and 5’- DNA ends (dT_n_-dT_n_ molecules) (Supplementary Table 1) in Buffer M275-10, 25°C. Example isotherms in which DNA was titrated into RecB^ΔNuc^CD are shown in Figure 2B and 2C, respectively. These isotherms show binding stoichiometries of two RecB^ΔNuc^CD molecules per DNA, consistent with the expectation that one RecB^ΔNuc^CD binds to each DNA end. RecB^ΔNuc^CD binding to a blunt ended DNA (Figure 2B) is associated with an unfavorable total enthalpy change (ΔH° > 0) suggesting that binding is dominated by the unfavorable enthalpic cost of melting the DNA base pairs, and that RecB^ΔNuc^CD binding to a blunt DNA end is driven by a favorable increase in entropy. On the other hand, RecB^ΔNuc^CD binding to the dT_30_-dT_30_ DNA molecule (Figure 2C) is associated with a large favorable enthalpy change (ΔH° < 0) due to the absence of the unfavorable enthalpy cost of DNA bp melting. The isotherms for other dT_n_dT_n_ molecules titrated into RecB^ΔNuc^CD are shown in Supplementary Figure 3.

The values of ΔH^°^ obtained from ITC experiments performed with a series of twin-tailed DNA molecules with ssDNA tail lengths between zero and 30 nucleotides are listed in Supplementary Table 3 and plotted in Figure 2D (solid blue dots). We are unable to obtain estimates of ΔG^°^ and TΔS^°^ because the binding affinities of RecB^ΔNuc^CD to all of these dT_n_-dT_n_ DNA molecules are too high to measure accurately by ITC (except for the blunt-ended dsDNA). As shown in Figure 2D and discussed above, the ΔH^°^ for RecB^ΔNuc^CD binding to a blunt DNA end is unfavorable (+10.4 ± 0.4 kcal/mol). However, ΔH^°^ for binding to all of the twin-tailed dT_n_-dT_n_ DNA ends is favorable and becomes more favorable with increasing tail length until reaching a plateau of ΔH^°^ = −51 ± 4 kcal/mol for n ≥ 15 (average of ΔH^°^ for dT_n_-dT_n_ with n=15, 20, 25, 30). For comparison, Figure 2D also shows the values of ΔH^°^ determined for RecBCD (red empty dots) and RecB^D1080A^CD (red solid dots), a nuclease deficient mutant of RecBCD, binding to the same DNA ends as determined previously^17^. For RecBCD, ΔH^°^ does not reach a plateau until n ≥ 20 and with a much more favorable ΔH^°^ = −75±2 kcal/mol.

The overall unfavorable ΔH^°^ = +10.4±0.4 kcal/mol for RecB^ΔNuc^CD binding to a blunt DNA end must be composed of unfavorable contributions from base pair melting^37–39^ and favorable contributions from protein-DNA interactions. The plateau value of ΔH^°^ = −51±4 kcal/mol for RecB^ΔNuc^CD provides an estimate of the favorable protein-DNA interactions in the absence of DNA bp melting (Eq. (2b). The ΔΔH^°^ = +61±4 kcal/mol, the difference between ΔH° for RecB^ΔNuc^CD binding to a blunt DNA end (n=0) and ΔH° for RecB^ΔNuc^CD binding to DNA with tails of n ≥ 15 (plateau value), provides an estimate for the unfavorable ΔH° contribution from bp melting (Eq. 3b). Based on available RecBCD-DNA structures^31^, a 3’-dT_20_ tail should be long enough to potentially reach the RecB^Nuc^ domain. This suggests that direct interactions between RecB^Nuc^ and the 3’-dT tail may be partially responsible for the more favorable ΔH^°^ for n > 15.

The much more favorable value of ΔH^°^ for RecBCD (−75±2 kcal/mol) vs. RecB^ΔNuc^CD (−51±4 kcal/mol) suggests a loss of protein-DNA interactions due to deletion of the RecB^Nuc^ domain. In addition, the value of ΔΔH^°^= 81±2 kcal/mol obtained for RecBCD^17^, which represents the contribution to ΔH° due to bp melting by RecBCD binding to a blunt DNA end, is also much larger than for RecB^ΔNuc^CD (61±4 kcal/mol). This difference of ∼20 kcal/mol suggests that RecB^ΔNuc^CD does not melt as many base pairs as RecBCD upon binding a blunt DNA.

### Cryo-EM experiments on RecB^ΔNuc^CD and RecB^ΔNuc^CD-DNA complexes

We performed cryo-EM experiments on RecB^ΔNuc^CD (Figure 3A) and RecB^ΔNuc^CD-DNA complexes formed by mixing RecB^ΔNuc^CD with an equal concentration of the 60 bp blunt-ended dsDNA (Figure 3B) in 20 mM Tris pH7.4, 50 mM NaCl and 4 mM MgCl_2_ with a final concentration of 0.025% amphipol added immediately before vitrification. This is the same buffer used by Wilkinson et al.^36^ and for our cryo-EM studies of RecBCD and RecBCD-DNA complexes^17^. We used the same analysis and classification methods as described for RecBCD^17^. 3D classification yielded only a single class for RecB^ΔNuc^CD consisting of 162,275 particles (Supplementary Figure 5). Increasing the number of classes or further sub-classification of this single class of RecB^ΔNuc^CD particles only identified an additional 1.5% of the particles as a class without defined features, and had no effect on the resulting 3D reconstruction of the RecB^ΔNuc^CD cryo-EM density map.

**Figure 3.**
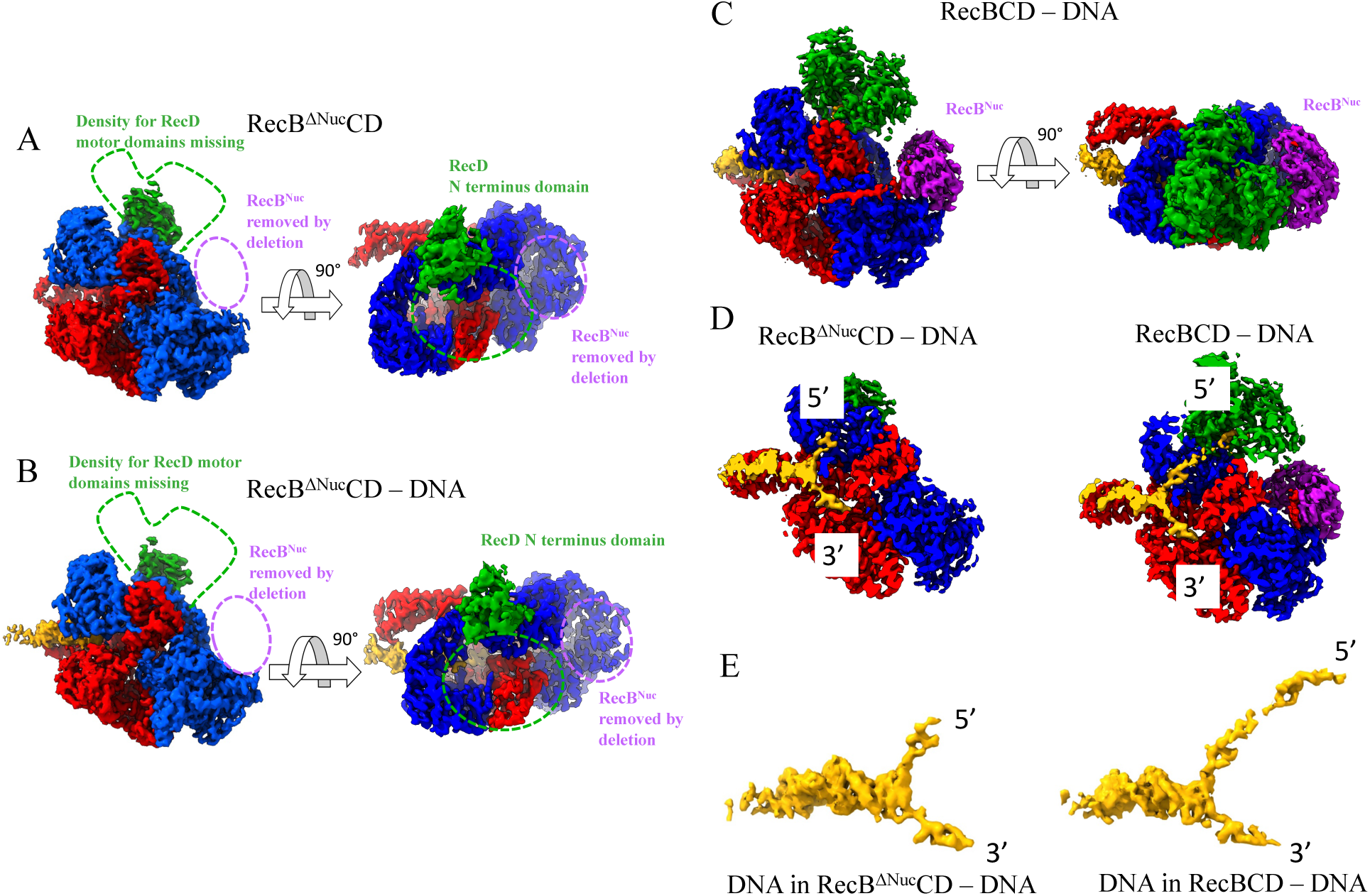
Structures of RecB^ΔNuc^CD and RecB^ΔNuc^CD bound to blunt-ended DNA. (A)- Cryo-EM structures with RecB in red, RecB^Nuc^ in purple, RecC in blue, dsDNA in yellow and RecD in green. (A)- RecB^ΔNuc^CD and (B)- RecB^ΔNuc^CD-DNA indicate weak map densities for most of RecD (dashed green lines). The position of the RecB^Nuc^ in the RecBCD structures is indicated by the dashed purple oval. (C)- Cryo-EM structure of RecBCD-DNA (showing RecB^Nuc^ docked) shows strong density for RecD. (D)- RecB^ΔNuc^CD-DNA structure shows only 4 base pairs melted from a blunt DNA end compared to 11 bp melted in the RecBCD-DNA structure (D). (E)- The cryo-EM maps from (D) showing only the DNA density.

We first used ITC to examine RecB^ΔNuc^CD binding to the 60 bp blunt-ended dsDNA under the cryo-EM solution conditions (20 mM Tris pH7.4, 50 mM NaCl and 4 mM MgCl_2_), which showed independent binding of RecB^ΔNuc^CD to both ends of the dsDNA molecule, with ΔH = +7.5±0.3 kcal/mol (open blue circle in Figure 2D, Supplementary Table 3). ITC experiments also showed stoichiometric binding of RecB^ΔNuc^CD to dsDNA possessing dT_20_-dT_20_ and dT_30_-dT_30_ ends with ΔH = −58±2 kcal/mol and ΔH = −56±3 kcal/mol, respectively (Figure 2D). This results in a ΔΔH° = ΔH_°melt_ = +65±4 kcal/mol, the same within error, as the value obtained in buffer M275-10 (Figure 2D, Supplementary Table 2). The cryo-EM experiments used a 1:1 molar ratio of [dsDNA] to [RecB^ΔNuc^CD], which is a two-fold excess of DNA ends over RecB^ΔNuc^CD. This should ensure that all RecB^ΔNuc^CD hetero-trimers are bound to DNA. The blunt ended DNA used in the cryo-EM experiment is the same as that used in the ITC studies reported here and in our previous studies of RecBCD^17^. Although using this DNA allows for comparison of the DNA binding studies for RecBCD and RecB^ΔNuc^CD, it prevents us from identifying the exact bases in the structure because the two DNA ends in the dsDNA have slightly different sequences. Analysis of the RecB^ΔNuc^CD- DNA complex data set yielded a major class (Class 1) of RecB^ΔNuc^CD-DNA consisting of 130,960 particles and a minor class of 26,084 particles of low resolution (Supplementary Figure 3B). No class representing DNA-free RecB^ΔNuc^CD was observed in the RecB^ΔNuc^CD-DNA data set.

Through further global refinement, we reconstructed cryo-EM density maps for RecB^ΔNuc^CD and RecB^ΔNuc^CD-DNA at 3.4 Å resolution (Figure 3A and 3B). (Figure 3A,B and Supplementary Figure 4). The 3D reconstruction for the minor class of RecB^ΔNuc^CD-DNA complexes resulted in an overall lower resolution of 7.2 Å that shows weak densities in many parts of the protein-DNA complex (data not shown). Attempts to classify RecB^ΔNuc^CD-DNA into more classes did not result in better cryo-EM density maps.

### RecD is disordered in RecB^ΔNuc^CD and RecB^ΔNuc^CD-DNA

The RecB^ΔNuc^CD particles show only a single structural class (Figure 3A, Supplementary Figure 5A) in stark contrast to the three structural classes previously observed for RecBCD particles^17^. The weak density corresponding to most of the RecD subunit suggests heterogeneity in the RecD subunit conformation within RecB^ΔNuc^CD. This weak density is not due to some sub-population in which RecD has dissociated since strong density corresponding to the N-terminus of RecD is evident (Figure 3A green). Results from sedimentation velocity experiments and denaturing gels also show the presence of all three subunits in our samples (Figure 1A and B).

Conformational heterogeneity of RecD is also evident in the RecB^ΔNuc^CD-DNA complex structure which showed weak density for most of the RecD subunit, except for the N-terminal domain (Figure 3B). This is in stark contrast to the RecBCD-DNA structures^17^, where 63.2% of the particles showed strong RecD density (Figure 3C and Hao et al.^17^). Our previous RecBCD study^17^ suggested that DNA binding eliminates the RecD conformational heterogeneity. This effect is not evident in the RecB^ΔNuc^CD-DNA structures based on the observation that no significant population shows RecB^ΔNuc^CD-DNA particles with full RecD density.

### RecB**^ΔNuc^**CD-DNA structures show fewer base pairs melted than RecBCD-DNA structures

Our thermodynamic studies suggest that RecB^ΔNuc^CD binding to blunt-ended DNA results in fewer bp melted compared to RecBCD-DNA binding. This conclusion is supported by the cryo-EM RecB^ΔNuc^CD-DNA structure (Figure 3D and 3E). The density map for the DNA in the RecB^ΔNuc^CD-DNA structure indicates that dsDNA melting does occur upon RecB^ΔNuc^CD binding to a blunt dsDNA end in the presence of Mg^2+^. However, only 3 unpaired nucleotides on the 5’ ended ssDNA and 4 unpaired nucleotides on the 3’ ended ssDNA are evident, indicating at least 4 base pairs are melted upon RecB^ΔNuc^CD binding to a blunt DNA end. This is significantly less than we observed in the major class (63.2%) of wild type RecBCD-DNA complexes, which showed melting of at least 11 base pairs using the same DNA substrate under the same conditions (Figure 3D right panel)^17^. The structure of RecB^ΔNuc^CD-DNA is similar to the minor (36.8%) RecBCD-DNA structural class observed in our previous study^17^, which also showed only 4 bp melted, along with weak RecD density and the absence of RecB^Nuc^ density. Taken together, the thermodynamic and structural studies presented here suggest that the RecB^Nuc^ domain plays a regulatory role in DNA bp melting. Furthermore, Fazio et al.^19^ demonstrated that RecB^ΔNuc^CD initiates DNA unwinding much slower than RecBCD and unwinds dsDNA at a slower rate than RecBCD indicating a regulatory role of the nuclease domain in DNA helicase activity.

### RecB^Nuc^ can exert long-range allosteric effects on DNA binding by affecting the conformations of RecB^2B^ and RecC^Cter^ domains

We next examined the structural differences between RecB^ΔNuc^CD and RecBCD^17^ in the DNA bound and DNA free states. We used the Matchmaker tool in UCSF Chimera^40^ to compare the structural models and carried out structural alignments on the full complexes rather than restricting the alignment to specific regions of the structures. The structures were matched on the most structurally similar chains as determined by the Needleman-Wunsch algorithm^41^. Local RMSD values were then calculated for different regions of the aligned structures as summarized in Table 1 and Supplementary Table 4. Supplementary Figure 4B shows colored maps detailing different regions of the RecBCD subunits to aid visualization of different domains in the protein complexes for subsequent structural comparisons.

**Table 1.**
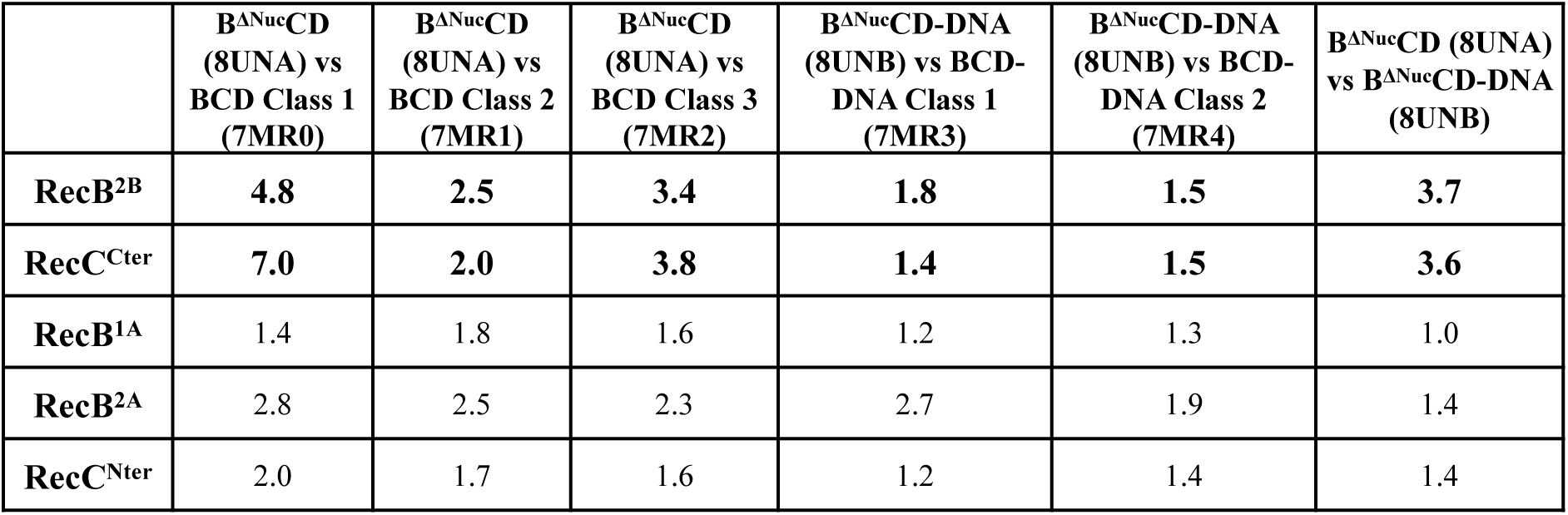
Local RMSD (Å) between various structures.

As displayed in Table 1, the 2B subdomain of RecB (RecB^2B^) and the C terminus of RecC (RecC^Cter^) show higher RMSD values when comparing certain structures. In contrast, consistently throughout our structural comparisons, the RecB motor domains (RecB^1A^, RecB^2A^) and the N terminal half of RecC (RecC^Nter^) show little conformational differences (Table 1).

The density maps of RecB^ΔNuc^CD (dark red, EMD-42396), and the major class (44.2%, EMD- 23952) of RecBCD (grey) where RecB^Nuc^ is docked, are aligned in Figure 4, with panels highlighting structural differences at specific regions of the complexes. These maps are displayed in Figure 4 from the same viewing angle as in Supplementary Figure 4A, which is color coded for different subunits in the protein complex to help with visualization. Differences in conformation in RecC^Cter^ (Figure 4A) and RecB^2B^ (Figure 4B) are highlighted. The RecC^Cter^ domains between these structures show an RMSD of 4.8 Å, while the RecB^2B^ domains show an RMSD of 7.0 Å (Table 1), both substantially higher than the overall resolution of these structures (3.7 Å for RecBCD EMD-23952 and 3.4 Å for RecB^ΔNuc^CD EMD-42396).

**Figure 4.**
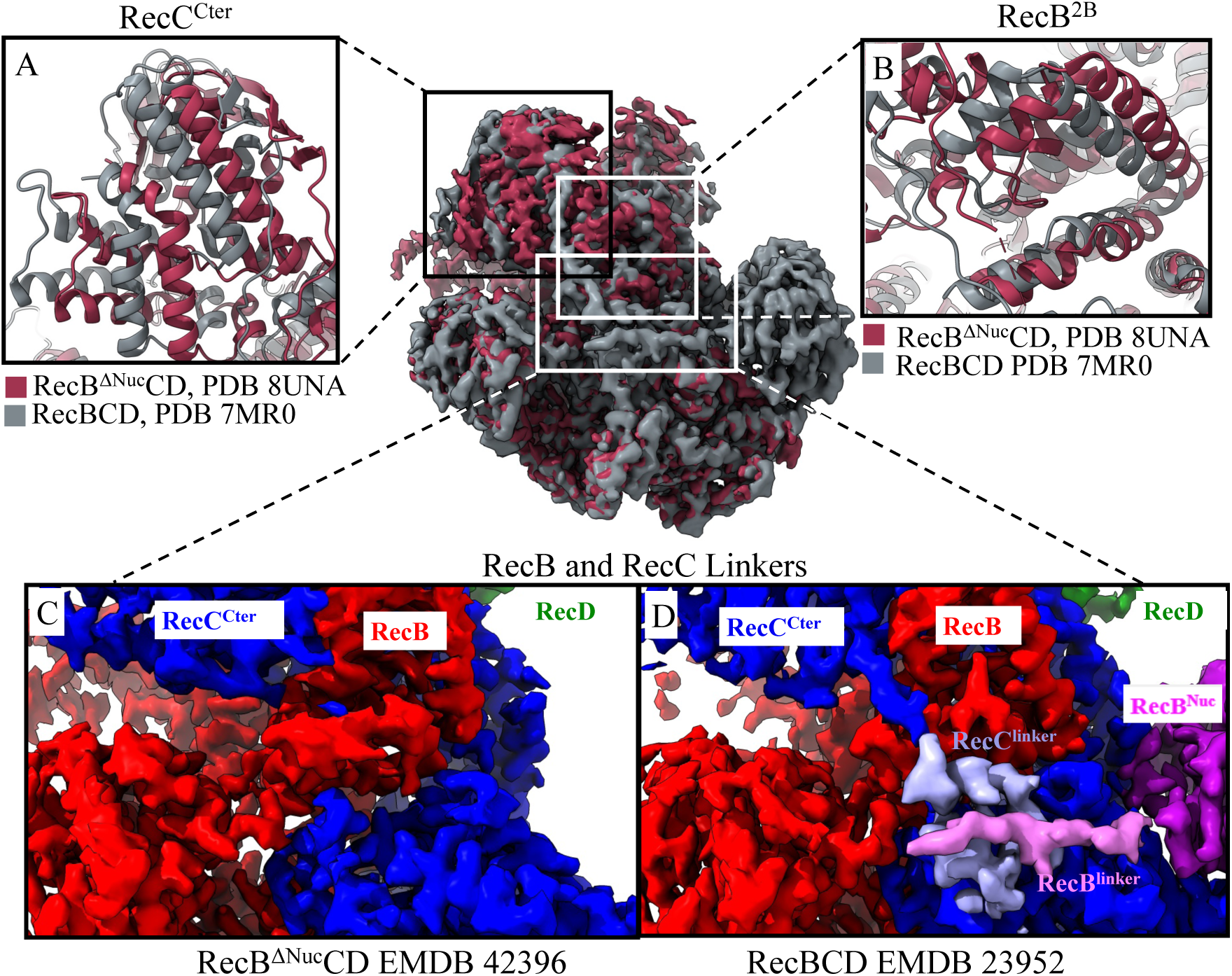
Comparison of RecB^ΔNuc^CD and RecBCD structures. Cryo-EM maps of RecB^ΔNuc^CD (dark red, EMD-42396) and RecBCD (grey, EMD-23952, with RecB^Nuc^ docked) were overlaid, and several regions of interest were compared in panels A-D. Panels A and B show structural shifts for RecC^Cter^ and RecB^2B^, respectively, between RecB^ΔNuc^CD and RecBCD. Panel C and Panel D show the map densities for RecB^Linker^ and RecC^Linker^. Densities for these regions are absent in RecB^ΔNuc^CD (C) but present in RecBCD (D).

However, upon comparing the RecB^ΔNuc^CD (8UNA) and RecBCD classes (7MR1 and 7MR2) where RecB^Nuc^ is undocked, the conformational differences in RecC^Cter^ and RecB^2B^ are much smaller (Table 1 and Supplementary Figure 7). This indicates that RecB^2B^ and RecC^Cter^ adopt similar conformations when RecB^Nuc^ is deleted or undocked. This suggests that the docking and undocking of RecB^Nuc^ can influence the conformations of RecC^Cter^ and RecB^2B^ in the DNA unbound state of RecBCD. It is possible that RecB^Nuc^ exerts long range allosteric effects on DNA binding by affecting conformations in RecB^2B^ and RecC^Cter^ which are in direct contact with DNA. RecB^Nuc^ is connected to the RecB motor domains via a long (∼60) amino acid linker, as shown in Supplementary Figure 4B. A similar linker also connects the N and C terminal domains of RecC (Supplementary Figure 4B). We note that the densities corresponding to the RecB^Linker^ (pink) and RecC^Linker^ (light blue) are clearly observed in RecBCD (EMD-23952, Figure 4D), but neither are observable in RecB^ΔNuc^CD (EMD-42396, Figure 4C), even though the RecB^Nuc^ deletion starts at amino acid 930, leaving the entire RecB^Linker^ intact. Given that the RecC^Linker^ is directly connected to the RecC^Cter^ domain, and that RecC^Cter^ and RecC^Linker^ interact with RecB^2B^, it is plausible that the docking and undocking of RecB^Nuc^ alters the interactions between the RecB^Linker^ and the RecC^Linker^, which may cause a shift in the conformations of RecB^2B^ and RecC^Cter^.

However, when comparing DNA bound structures, RecBCD-DNA (with RecB^Nuc^ docked, EMD- 23952) and RecB^ΔNuc^CD-DNA (EMD-42397), no conformation differences are observed for RecB^2B^ and RecC^Cter^, despite the differences in RecB^Nuc^ docking and the RecB^Linker^ and RecC^Linker^ densities. A comparison of these structures is shown in Figure 5 and summarized in Table 1. In addition to this, we note that RecB^2B^ (and RecC^Cter^) shows the same conformations between the two classes of RecBCD-DNA structures, where one has RecB^Nuc^ docked and the other undocked (Supplementary 4). These observations indicate that the docking and undocking of RecB^Nuc^ does not affect the conformations of RecB^2B^ and RecC^Cter^ in DNA bound RecBCD.

**Figure 5.**
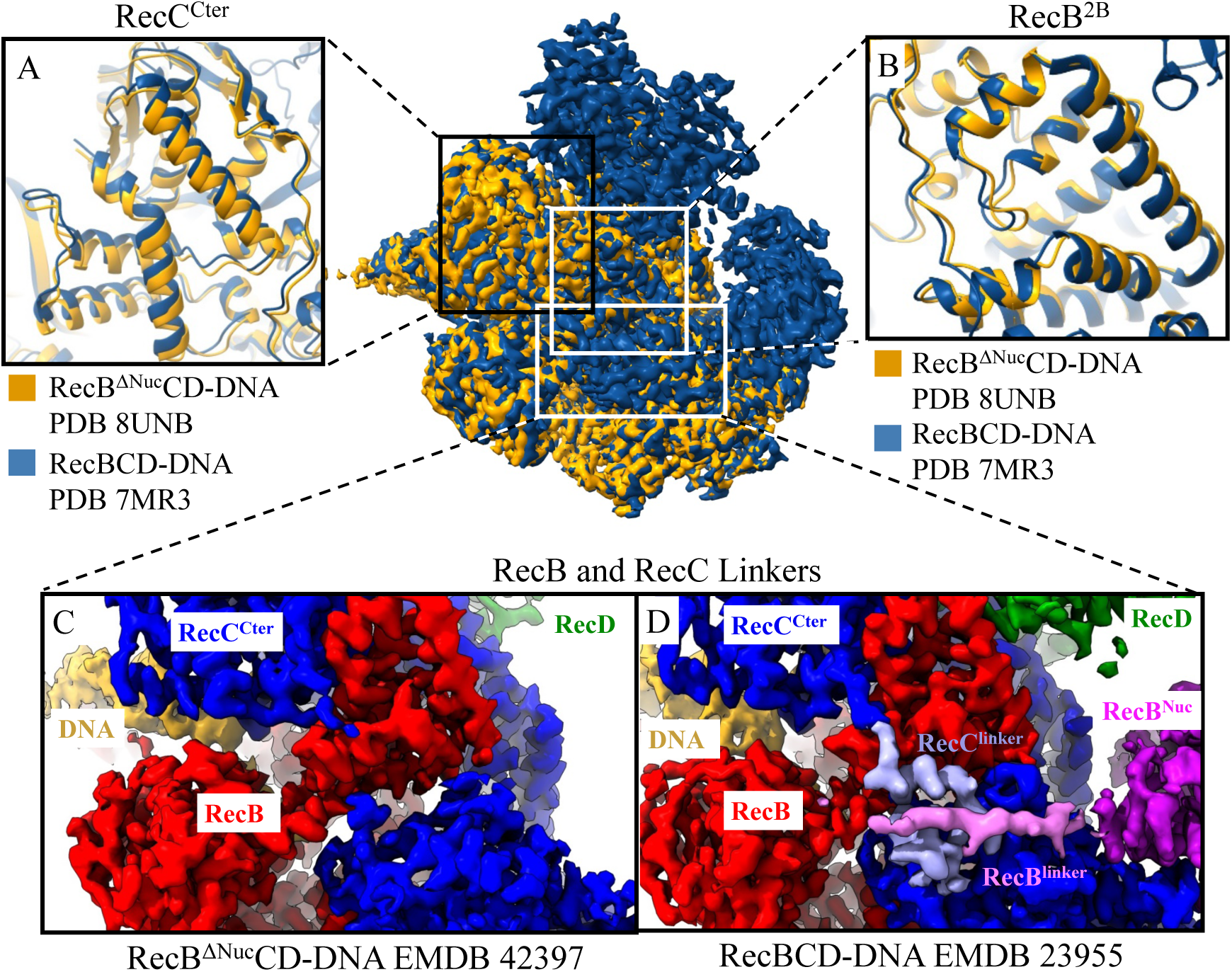
Comparison of RecB^ΔNuc^CD-DNA and RecBCD-DNA structures. Cryo-EM maps of RecB^ΔNuc^CD-DNA (yellow) and RecBCD-DNA (dark blue, 7MR3, with RecB^Nuc^ docked) are overlaid and several regions of interest are compared in panels A-D. Panels A and B show the structural similarities between RecC^Cter^ and RecB^2B^ domains, respectively. Panels C and D show that the map densities corresponding to RecC^Linker^ and RecB^Linker^ are missing in RecB^ΔNuc^CD-DNA but present in RecBCD-DNA.

We also note that DNA binding to RecB^ΔNuc^CD causes conformational shifts in RecB^2B^ and RecC^Cter^. Supplementary Figure 7 compares the density maps of RecB^ΔNuc^CD-DNA (EMD-42397) and RecB^ΔNuc^CD (EMD-42396) showing the conformational differences in RecB^2B^ (Supplementary Figure 7A) and RecC^Cter^ (Supplementary Figure 7B). Structural comparisons between the various classes of RecBCD and RecBCD-DNA are shown in Supplementary Figure 9 and 10, and summarized in Supplementary Table 4. The RMSD values (of RecB^2B^ and RecC^Cter^) in these comparisons are consistently higher than the RecB motor domains and RecC^Nter^ RMSDs, suggesting that DNA binding affects the RecB^2B^ and RecC^Cter^ conformational states in RecBCD as well. Based on these comparisons, we hypothesize that the docking/undocking (or deletion) of RecB^Nuc^ primarily influences the DNA free state of RecBCD but not the DNA bound state by regulating the conformational states of the RecB^2B^ and RecC^Cter^ domains through RecB^Linker^-RecC^Linker^ interactions.

## Discussion

RecBCD functions in repair of double stranded DNA breaks by binding to a dsDNA end and processively unwinding the DNA duplex while degrading the resulting ssDNA using a single nuclease active site on RecB^Nuc^ ^20–22^. Upon recognizing a Chi site, the 3’ ssDNA end becomes protected from further degradation and RecBCD facilitates the loading of a RecA filament onto the 3’ ended ssDNA that is used to initiate a homologous recombination event and subsequent DNA repair process^22–27^. The facilitation of RecA loading is thought to involve direct interactions of RecA with the nuclease domain (RecB^Nuc)^. Structural data suggest that RecB^Nuc^ must either undergo a substantial conformational change^42^ or become undocked from its location in the crystal structure in order to interact with RecA^28^.

We previously demonstrated that the RecB^Nuc^ within RecBCD and the RecBCD-DNA complex appears to be dynamic, even in the absence of Chi recognition^17^. Significant populations of both RecBCD and RecBCD-DNA complexes show structures in which RecB^Nuc^ is not visible, suggesting that it has become undocked, but still covalently bound via its flexible linker. More recent studies demonstrated that removal of RecB^Nuc^ directly impacts the initiation of dsDNA unwinding, suggesting an allosteric role of RecB^Nuc^ in regulating motor engagement with the DNA ends^13,19^. Using a combined structural and thermodynamic approach, we show here that RecB^Nuc^ also influences RecBCD binding affinity to DNA ends and the extent of base pair melting upon binding to a blunt DNA end. We also suggest structural explanations for how RecB^Nuc^ might play an allosteric role in regulating RecBCD-DNA interactions.

### RecB^Nuc^ regulates RecBCD-DNA binding and base pair melting

Under the solution conditions used in our experiments, we find that RecB^ΔNuc^CD binds with higher affinity than RecBCD to all DNA ends examined. RecBCD and RecB^ΔNuc^CD bind to blunt ended DNA with similar binding constants at 275 mM NaCl (Buffer M275-10, 25°C) (K_BΔNucCD_=1.2 (±0.4)×10^7^M^-1^, Supplementary Table 2; K_BCD_=7.0±0.8 x 10^6^ M^-1^ Hao et al.^17^). However, the difference in affinity becomes greater for ends with ssDNA flanking regions. Furthermore, both RecBCD and RecB^ΔNuc^CD binding to a blunt DNA end are enthalpically unfavorable, although with different values of ΔH°, with ΔH° = +4.6±0.2 kcal/mol for RecBCD and ΔH° = +10.4±0.4 kcal/mol for RecB^ΔNuc^CD. Hence, the binding constants, and thus ΔG° for the two enzymes will differ more at a different temperature. Although we can accurately measure the ΔH° for binding of RecBCD and RecB^ΔNuc^CD to DNA ends possessing twin ssDNA tails, the binding constants are too high to measure in the solution conditions that we used.

RecB^ΔNuc^CD binding to a blunt DNA end is associated with a more unfavorable enthalpy change (ΔH_obs_ = +10.4 (±0.4) kcal/mol) than for RecBCD (ΔH_obs_ = +4.6 (±0.2) kcal/mol) (Buffer M275-10, 25 °C). This positive enthalpy change reflects the favorable interactions (negative enthalpy) between the protein and DNA, and the unfavorable (positive enthalpy) melting of dsDNA base pairs. A DNA end with unpaired ssDNA tails (pre-melted) removes the enthalpic cost of DNA base pair melting. Binding of RecB^ΔNuc^CD to a DNA end with sufficiently long unpaired ssDNA tails should only show contributions from the favorable protein-DNA interactions. Comparisons of RecB^ΔNuc^CD binding to a blunt DNA end vs a fully pre-melted DNA (average of dT_15_/dT_15_ through dT_30_/dT_30_) yields a ΔΔH° = +61 ±4 kcal/mol which provides an estimate of the cost of DNA melting by RecB^ΔNuc^CD. Using the same approach, RecBCD has a ΔΔH° = +81 ±2 kcal/mol and RecBC has ΔΔH° = +47 ±7 kcal/mol^17,34^. The ΔΔH° for RecB^ΔNuc^CD is larger than measured for RecBC but smaller than for RecBCD, suggesting that RecB^ΔNuc^CD is able to melt more base pairs from a blunt DNA end than RecBC but less than RecBCD.

The favorable ΔH° for RecB^ΔNuc^CD binding to DNA ends increases (ΔH° decreases) as the ssDNA tail length increases, reaching a plateau at n =15 nucleotides. For RecBCD, the plateau occurs at n=17-18 nucleotides (Figure 2D^17^). This suggests that RecB^ΔNuc^CD and RecBCD can melt as many as 15 bp, or 17- 18 bp, respectively. The resulting enthalpic cost per bp would be ΔH° = +4.1 ±0.2 kcal/mol bp for RecB^ΔNuc^CD and ΔH° = +4.5 ±0.3 kcal/mol bp for RecBCD. These estimates are in line with previously published enthalpic cost of DNA melting^43–45^. For comparison, Wong and Lohman^34^ demonstrated that RecBC melts ∼6 bp from a blunt DNA end. This results in a ΔH = +8 ±1 kcal/mol bp, based on the ΔΔH° = +47 ±7 kcal/mol. This estimate is also in line with various reports estimating the enthalpic cost of DNA base pair melting^43–45^. Based on this estimate and the ΔΔH° for RecBCD and RecB^ΔNuc^CD, RecBCD is predicted to melt 9-12 bp^17^ and RecB^ΔNuc^CD is predicted to melt 7-9 bp.

The cryo-EM structures of DNA complexes with RecBCD^17^ and RecB^ΔNuc^CD presented in this study are consistent with our thermodynamic measurements. The RecB^ΔNuc^CD-DNA complexes showed a single class of structures with map density corresponding to 4 bp melted from a blunt dsDNA end. This is in stark contrast to the heterogeneity of RecBCD-DNA complexes observed previously^17^, where two substantial populations of RecBCD-DNA complexes were identified: a major class showed clear density corresponding to melting of at least 11 bp from a blunt DNA end, and a minor class showing melting of only 4 bp. The class of RecBCD-DNA structures showing melting of 11 bp also showed clear density for the RecB nuclease domain, whereas density for the RecB nuclease domain was not observed for the class of RecBCD-DNA structures showing melting of 4 bp. The structures of RecB^ΔNuc^CD-DNA and the minor class of RecBCD-DNA show obvious similarities: 1) both show the absence of density for the RecB^Nuc^; 2) a largely flexible RecD subunit; and 3) density corresponding to melting of only 4 bp of DNA.

It is important to note that ensemble DNA unwinding experiments showed that only about 80% of RecBCD initiates DNA unwinding rapidly from a blunt DNA end^46^. This could mean that the ∼37% of RecBCD-DNA complexes showing only melting of 4 bp of DNA represents a complex that is unable to rapidly initiate DNA unwinding. Recent studies also demonstrated that RecB^ΔNuc^CD unwinds DNA at a slower rate than RecBCD^13,19^. Furthermore, RecB^ΔNuc^CD initiates DNA unwinding poorly from a blunt DNA end compared to RecBCD, but a DNA end possessing a 5’-dT_10_ flanking ssDNA enhances initiation of DNA unwinding by RecB^ΔNuc^CD^19^. Together, these results suggest that RecB^Nuc^ and its proper docking is required for DNA melting by RecBCD and RecD engagement with 5’ ssDNA tail is required for proper initiation of DNA unwinding.

### RecB^Nuc^ may exert an allosteric effect through RecB^linker^-RecC^linker^ interactions

Our results suggest RecB^Nuc^ exerts an allosteric effect on RecBCD binding and bp melting of dsDNA ends^19,47^. This is unexpected because RecB^Nuc^ is located far from potential direct interactions with DNA in all structural studies in which RecB^Nuc^ is observed. How might RecB^Nuc^ achieve such long-range allosteric effects? Comparisons of our current and previous^17^ cryo-EM structures show that when RecB^Nuc^ density is not observed (RecB^Nuc^ is either undocked or deleted), the density corresponding to RecB^Linker^ and RecC^Linker^ are also not observed. Structural comparisons also indicate significant conformational variability for the RecB^2B^ and RecC^Cter^ domains within the apo form of RecBCD. This variability correlates with whether the RecB^Nuc^ domain is docked or undocked. However, in the RecB^ΔNuc^CD structure without DNA binding, the conformation of RecB^2B^ and RecC^Cter^ are much less variable. Given that RecC^Linker^ not only connects RecC^Cter^ to the rest of RecC but also interacts with RecB^2B^, we suggest that RecB^Nuc^ may allosterically regulate RecBCD-DNA interactions through linker-linker interactions between RecB and RecC.

The RecBCD structure involves direct interactions between the 2B sub-domains of RecB and RecC that stabilize the hetero-dimer^48^. Dimerization of the 2B sub-domains of other SF1A helicases, such as PcrA^49^, Rep^50^, *E. coli* UvrD^51^, and *M. tuberculosis* UvrD1^52,53^ also play a role in activation of the helicase by dimerization. The 2B sub-domains in the monomeric forms of these enzymes are auto-inhibitory for monomer helicase activity^54^ due to direct interactions of the 2B sub-domain with duplex DNA^49,51,53^. This auto-inhibition is removed via dimerization that occurs between the 2B sub-domains of the two subunits. Dimerization results in a large rotational movement of the 2B sub-domain that prevents its interaction with the duplex DNA, thus relieving the auto-inhibition^53^. Thus the 2B domains play regulatory roles in many of these SF1A helicases.

#### Material and Methods

**Buffers**

Reagent grade chemicals and double-distilled water further deionized with a Milli-Q purification system (Millipore Corp., Bedford, MA) were used to make all buffers. Buffer A is 50 mM Tris HCl, pH 7.5, 10% sucrose. Buffer C is 20 mM potassium phosphate, pH 6.8, 0.1 mM 2-mercaptoethanol, 0.1mM EDTA, 10% (v/v) glycerol. Buffer M contains 20 mM MOPS-NaOH, pH 7.0, 1mM 2-mercaptoethanol, 5% (v/v) glycerol. Stock concentration of MgCl_2_ solutions was determined by measuring the refractive index using a Mark II refractometer (Leica Inc., Buffalo, NY). All DNA binding experiments were performed in Buffer M or cryo-EM Buffer (20 mM Tris pH 7.4, 50 mM NaCl, 4 mM MgCl_2_). Buffer nomenclature is Buffer MX-Y and TX-Y, where X indicates the [NaCl] and Y indicates the [MgCl_2_] in mM concentration units (e.g., Buffer M30-10 has 30 mM NaCl and 10 mM MgCl_2_).

### Proteins and DNA

RecB^ΔNuc^CD hetero-trimer was purified as described previously for RecBCD^46,47,55^. Purified RecB^ΔNuc^CD was dialyzed into Buffer C, aliquoted, flash-frozen in liquid nitrogen and stored at −80°C. RecB^ΔNuc^CD concentration was determined by absorbance in Buffer C, using an extinction coefficient^47^ of ε_280_=4.11×10^5^ M^-1^ cm^-1^. Bovine serum albumin (BSA, from Sigma St. Louis, MO) concentration was determined by absorbance using an extinction coefficient of ε_280_=4.3×10^4^M^-1^cm^-1^ in Buffer C ^16^.

Oligodeoxynucleotides were synthesized using a MerMade 4 synthesizer (Bioautomation, Plano, TX) with phosphonamidite reagents (Glen Research, Sterling, VA) and purified as described^56^. Concentrations of each oligodeoxynucleotide were determined spectrophotometrically as described^44,47^. Double stranded DNA was formed by annealing the two complementary single stranded oligodeoxynucleotides by heating the mixture to 95°C for 5 minutes followed by slow cooling to 25°C.

### Sedimentation Velocity

Sedimentation velocity experiments were performed at 42000 rpm, 25°C, using an An50Ti rotor in an Optima XL-A analytical ultracentrifuge (Beckman Coulter, Fullerton, CA, USA). The concentrations of RecB^ΔNuc^CD used were between 0.3-1 µM, and sedimentation was monitored by absorbance at 280 nm. SEDNTERP [44] was used to determine the density and viscosity of the buffers at 20°C and the partial specific volume of RecB^ΔNuc^CD (0.736ml/g) in Buffer M50-10. Sedimentation data were analyzed using SEDFIT to yield continuous sedimentation coefficient distributions, c(s)^57,58^. The sedimentation coefficients were converted to s_20,w_ using SEDFIT^57,58^ and plotted in Figure 1B.

### Fluorescence Titrations to determine Equilibrium Binding constants

Direct titrations of the Cy3 labeled reference DNA by RecB^ΔNuc^CD were performed and analyzed to obtain equilibrium binding constants as described for RecBCD^16^. The equilibrium binding constants for RecB^ΔNuc^CD binding to unlabeled DNA were determined by a competition approach as described for RecBCD^16^.

### Isothermal Titration Calorimetry

ITC experiments were performed using a VP-ITC calorimeter (Malvern Panalytical, Malvern, UK) as described^16,59^. Solutions of RecB^ΔNuc^CD and DNA were extensively dialyzed against the reaction buffer at 4°C. Samples were then centrifuged to remove any insoluble particulates and degassed before use. A solution of RecB^ΔNuc^CD (0.3 to 1µM in the sample cell) was titrated with 10 µL injections of DNA (3-5µM in the syringe) at 5 min intervals with a stirring rate of 130 rpm. Separate control experiments were performed to determine the heat of dilution for each injection by injecting the same volumes of DNA into the sample cell containing only buffer. An N independent and identical sites model was used to analyze the total heat after the *i*-th injection (*Q_i_ ^tot^*) as a function of [DNA] using Eq. (4) to obtain the observed enthalpy change (ΔH_obs_) and equilibrium binding constant (K_obs_) for RecB^ΔNuc^CD binding to each DNA end, and the binding stoichiometry (N), although N was floated, the non-linear least squares analysis always yielded N=2 within a 2% uncertainty.

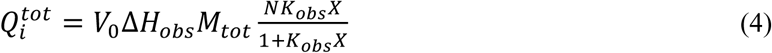

We emphasize that in equation (4) ΔH_obs_ and K_obs_ are the values for RecB^ΔNuc^CD binding to one end of a dsDNA substrate. V_0_ is the volume of the calorimetry cell (1.43ml), M_tot_ is the total DNA concentration and X is the free RecB^ΔNuc^CD concentration and is obtained by solving Eq. (5),

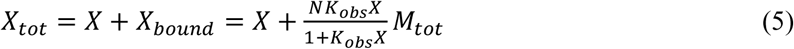

where X_tot_ is the total RecB^ΔNuc^CD concentration of in the cell after the *i*^th^ injection. When K_obs_ can be measured (10^3^M^-1^<K_obs_<10^9^M^-1^), the (1 M) standard state binding free energy (ΔG°) and the entropy change of binding (TΔS°) are calculated from eqs. (6) and (7), respectively. All uncertainties are reported

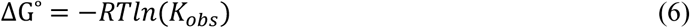

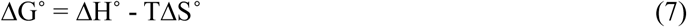

at the 68% confidence limit (± one standard deviation).

### Cryo-EM sample preparation and imaging

Cryo-EM sample preparation and imaging was carried out as described in our previous study^17^. For preparation of cryo-EM grids, RecB^ΔNuc^CD was extensively dialyzed vs. a buffer containing 20mM Tris, pH 7.4, 50mM NaCl and 4mM MgCl_2_. RecB^ΔNuc^CD was concentrated to 10µM (determined spectrophotometrically) using Vivaspin 500 centrifugal concentrators (Sartorius Stedim Biotech, NY) followed by centrifugation to remove any insoluble material. RecB^ΔNuc^CD-DNA complexes were formed by adding the blunt-ended dsDNA substrate (in the same buffer as RecB^ΔNuc^CD) to a final concentration of 15µM and allowed to incubate on ice for at least 15 minutes. Right before grid preparation, amphipol A8- 35 (Anatrace, OH) was added to a final concentration of 0.025% for both RecB^ΔNuc^CD and RecB^ΔNuc^CD samples.

Grids were prepared and imaged at the Washington University Center for Cellular Imaging (WUCCI). Immediately after addition of amphipol, 3µl of RecB^ΔNuc^CD or RecB^ΔNuc^CD solution was applied to holey carbon grids (Quantifoil R2/2 300mesh) that were glow discharged. The grids were blotted using FEI Vitrobot Mark IV (FEI) at 100% humidity for 2s and plunge-frozen into liquid ethane. The prepared grids were imaged using a Titan Krios (FEI) G3 electron microscope operating at 300kV. The cryo-EM data were recorded in counting mode on a K2-Summit direct electron detector (Gatan) with a GIF Quantum energy filter at slit width 20 eV. Images were acquired with EPU software (ThermoFisher Scientific) with a pixel size of 1.1 Å and a defocus range of −1.0 to −2.5µm. Data were collected with a dose rate of 1.65 e/Å^2^ per frame over a total of 40 frames with a frame rate of 0.2 s/frame and a total dose of 66 e/Å^2^.

### Image processing and model building

The image processing workflows are summarized in Supplementary Figure 3 and follow the same steps as in our previous study^17^. The RecB^ΔNuc^CD and RecB^ΔNuc^CD-DNA datasets were processed similarly with similar strategies. Corrections for beam-induced motion and dose weighting were performed using MotionCorr2^60^. The contrast transfer function (CTF) was determined using GCTF^61^. Gautomatch was used for automated particle picking. Extracted particles were subjected to two rounds of two-dimensional (2D) classification using a particle box size of 250 pixels. The resulting particles were used to generate a de novo three-dimensional (3D) initial model using Relion 3^62^. 3D classifications were carried out using the initial model as a reference map. For RecB^ΔNuc^CD, 3D classification produced one structural class of RecB^ΔNuc^CD particles with 162275 particles. The class lacking defined features was discarded. 3D refinement was carried out, resulting in an overall resolution of 3.4Å (Supplementary Figure 3A and Supplementary Figure 4B). For RecB^ΔNuc^CD-DNA, 3D classification produced one class of high quality RecB^ΔNuc^CD-DNA (130960 particles, Supplementary Figure 3B). 3D refinement produced maps with overall resolution of 3.4Å (Supplementary Figure 4E). Relion 3^62^ was used to calculate local resolutions.

For model building, the atomic model of RecBCD and RecBCD in complex with dsDNA from our previous cryo-EM study (PDB codes 7MR0 and 7MR3)^17^ was used as a template for both our RecB^ΔNuc^CD and RecB^ΔNuc^CD-DNA structures. It was first fit into our cryo-EM maps using UCSF Chimera^40^. An initial round of rigid body refinement was performed using PHENIX^64^, followed by cycles of real_space_refine in PHENIX^64^ and manual model building in COOT^65^. Structural figures were made using UCSF ChimeraX^66^. Model statistics (Supplementary Tables 4) were generated using PHENIX real-space refinement^64^. The final cryo-EM maps of RecB^ΔNuc^CD and RecB^ΔNuc^CD-DNA have been deposited to the Electron Microscopy Data Bank with codes EMD-42396 and EMD-42397, respectively. Corresponding atomic models have been deposited in the Protein Data Bank with accession codes 8UNA and 8UNB.

## Accession numbers

For RecB^ΔNuc^CD, PDB: 8UNA, EMBD: 42396

For RecB^ΔNuc^CD-DNA PDB: 8UNB, EMBD: 42397

See also Supplementary Table 4.

## Acknowledgements

We thank Alex Kozlov for help with the ITC experiments and for comments on the manuscript, Nicole Fazio for helpful discussions, Thang Ho for DNA synthesis and purification, and Michael Rau (WUCCI) for help in preparing and imaging the cryo-EM grids. This research was supported in part by NIH grants R01GM45948 and R35GM136632 (to TML) and R01GM138854 (to RZ).

## Supplementary Information

**Supplementary Table 1.**
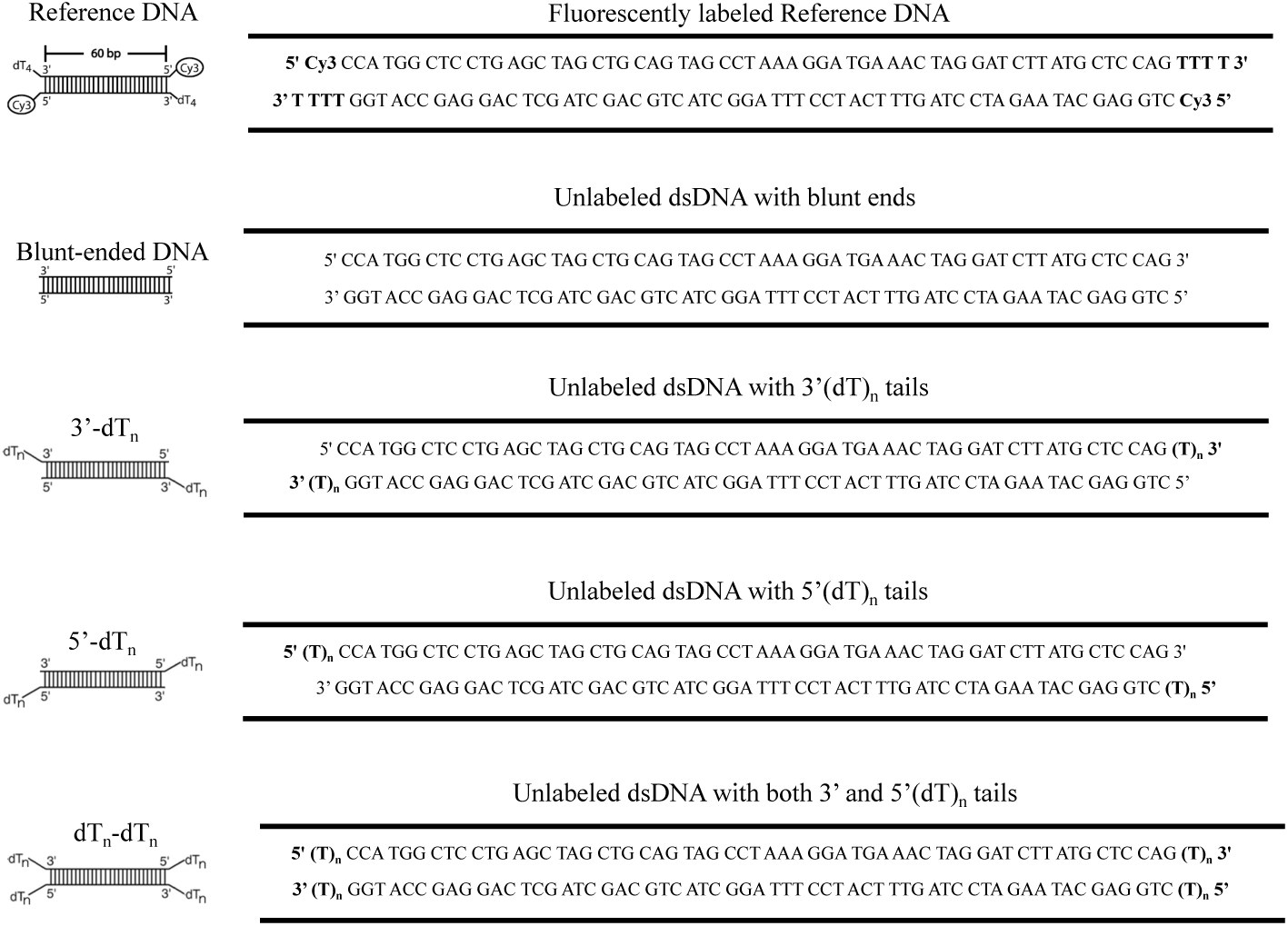
Sequences of DNA Substrates.

**Supplementary Table 2.** Equilibrium constants of RecBCD (K_BCD_), RecB^ΔNuc^CD (K_BΔNucCD_) binding to 3’-dT_n_ or 5’-dT_n_ DNA ends in Buffer M275-10, 25°C, from fluorescence titrations.

| Substrates | $K_{B\Delta NucCD}$ ($10^8 M^{-1}$) | $K_{BCD}$ ($10^8 M^{-1}$) |
| --- | --- | --- |
| Blunt Ended | 0.47±0.04 | 0.14±0.05 |
| 3'-dT<sub>2</sub> | 0.96±0.06 | 0.21±0.06 |
| 3'-dT<sub>4</sub> | 3.3±0.2 | 0.47±0.02 |
| 3'-dT<sub>6</sub> | 9.5±0.5 | 1.1±0.1 |
| 3'-dT<sub>8</sub> | 7.2±0.3 |  |
| 3'-dT<sub>10</sub> | 5.9±0.6 | 0.5±0.1 |
| 3'-dT<sub>15</sub> | 0.6±0.1 | 0.1±0.01 |
| 5'-dT<sub>6</sub> | 1.4±0.1 | 1.1±0.1 |
| 5'-dT<sub>8</sub> | 2.3±0.1 |  |
| 5'-dT<sub>10</sub> | 6.4±0.6 | 2.2±0.4 |
| 5'-dT<sub>15</sub> | 6.5±0.7 | 2.3±0.2 |

**Supplementary Table 3.**
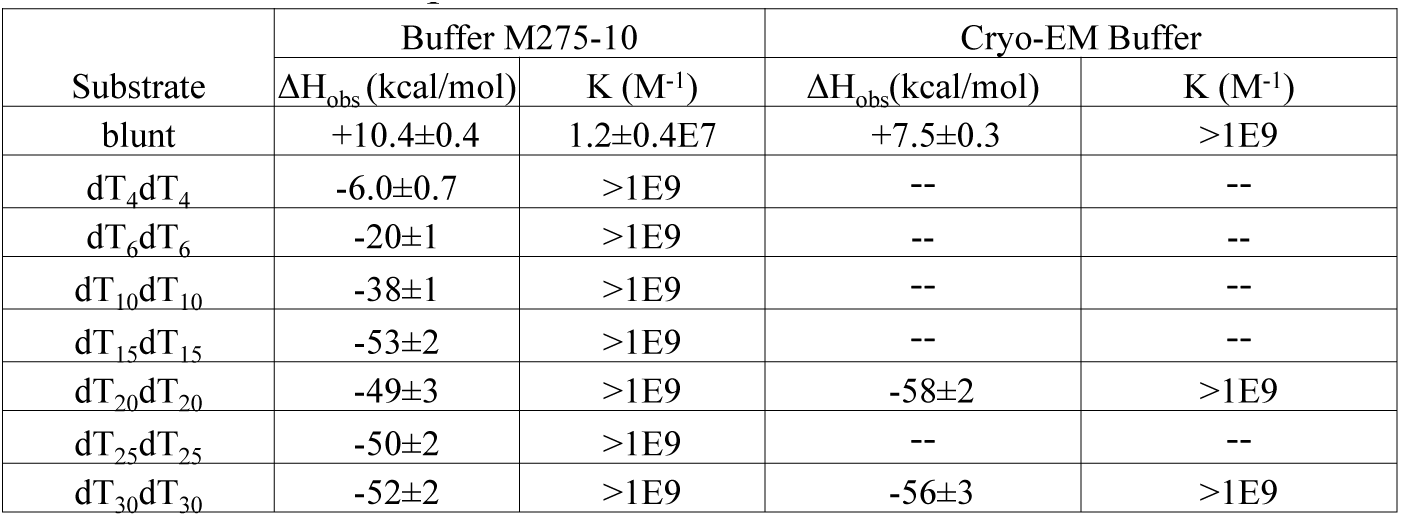
Parameters of ITC experiments for RecB^ΔNuc^CD.

| Substrate | Buffer M275-10 |  | Cryo-EM Buffer |  |
| --- | --- | --- | --- | --- |
| | $\Delta H_{\text{obs}}$ (kcal/mol) | K (M<sup>-1</sup>) | $\Delta H_{\text{obs}}$ (kcal/mol) | K (M<sup>-1</sup>) |
| blunt | +10.4±0.4 | 1.2±0.4E7 | +7.5±0.3 | >1E9 |
| dT<sub>4</sub>dT<sub>4</sub> | -6.0±0.7 | >1E9 | -- | -- |
| dT<sub>6</sub>dT<sub>6</sub> | -20±1 | >1E9 | -- | -- |
| dT<sub>10</sub>dT<sub>10</sub> | -38±1 | >1E9 | -- | -- |
| dT<sub>15</sub>dT<sub>15</sub> | -53±2 | >1E9 | -- | -- |
| dT<sub>20</sub>dT<sub>20</sub> | -49±3 | >1E9 | -58±2 | >1E9 |
| dT<sub>25</sub>dT<sub>25</sub> | -50±2 | >1E9 | -- | -- |
| dT<sub>30</sub>dT<sub>30</sub> | -52±2 | >1E9 | -56±3 | >1E9 |

**Supplementary Table 4.**
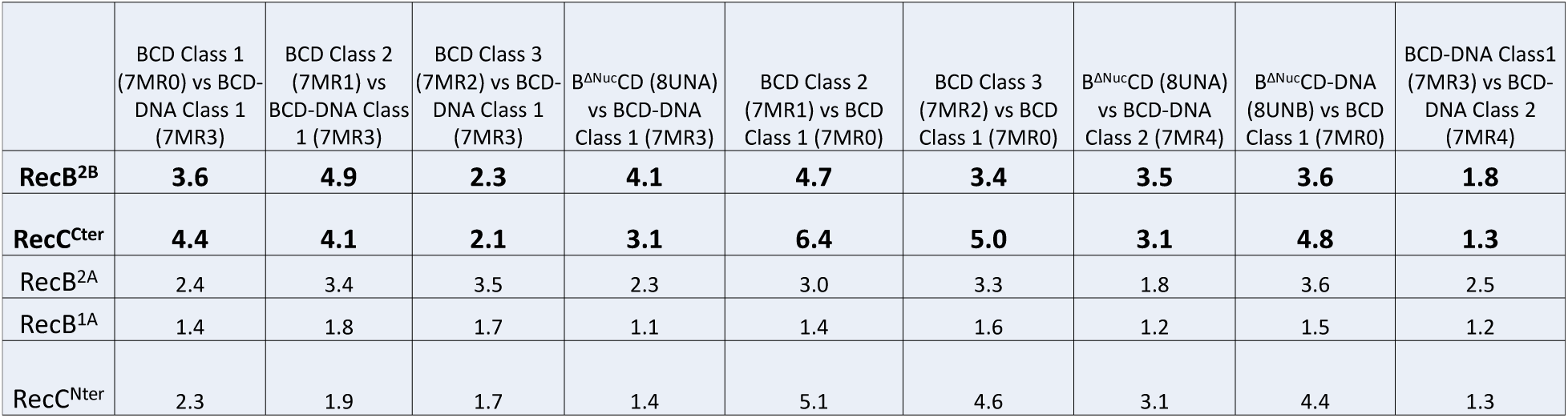
Local RMSD (Å) between regions of various structures.

**Supplementary Table 5.** Cryo-EM statistics for RecB^ΔNuc^CD and RecB^ΔNuc^CD-DNA structures.

| **Models** | **B<sup>ΔNuc</sup>CD** | **B<sup>ΔNuc</sup>CD-DNA** |
| --- | --- | --- |
| **PDB Accession Code** | **8UNA** | **8UNB** |
| **Model Composition** |  |  |
| Nonhydrogen Atoms | 16387 | 17075 |
| Protein Residues | 2036 | 2038 |
| Nucleotides | 0 | 34 |
| **Bonds (RMSD)** |  |  |
| Length (Å) | 0.004 | 0.002 |
| Angle (°) | 0.820 | 0.480 |
| **Validation** |  |  |
| MoProbiy Score | 1.54 | 1.63 |
| Clash Score | 4.83 | 7.17 |
| Rotamer outliers (%) | 0.17 | 0 |
| Cβ outliers (%) | 0 | 0 |
| CaBLAM outliers (%) | 2.26 | 2.23 |
| **Ramachandran plot (%)** |  |  |
| Outliers | 0.05 | 0.05 |
| Allowed | 4.16 | 3.50 |
| Favored | 95.79 | 96.45 |
| **B-factors (Å<sup>2</sup>)** |  |  |
| Protein | 56.79 | 66.17 |
| Nucleotides |  | 176.59 |

**Supplementary Figure 1.**
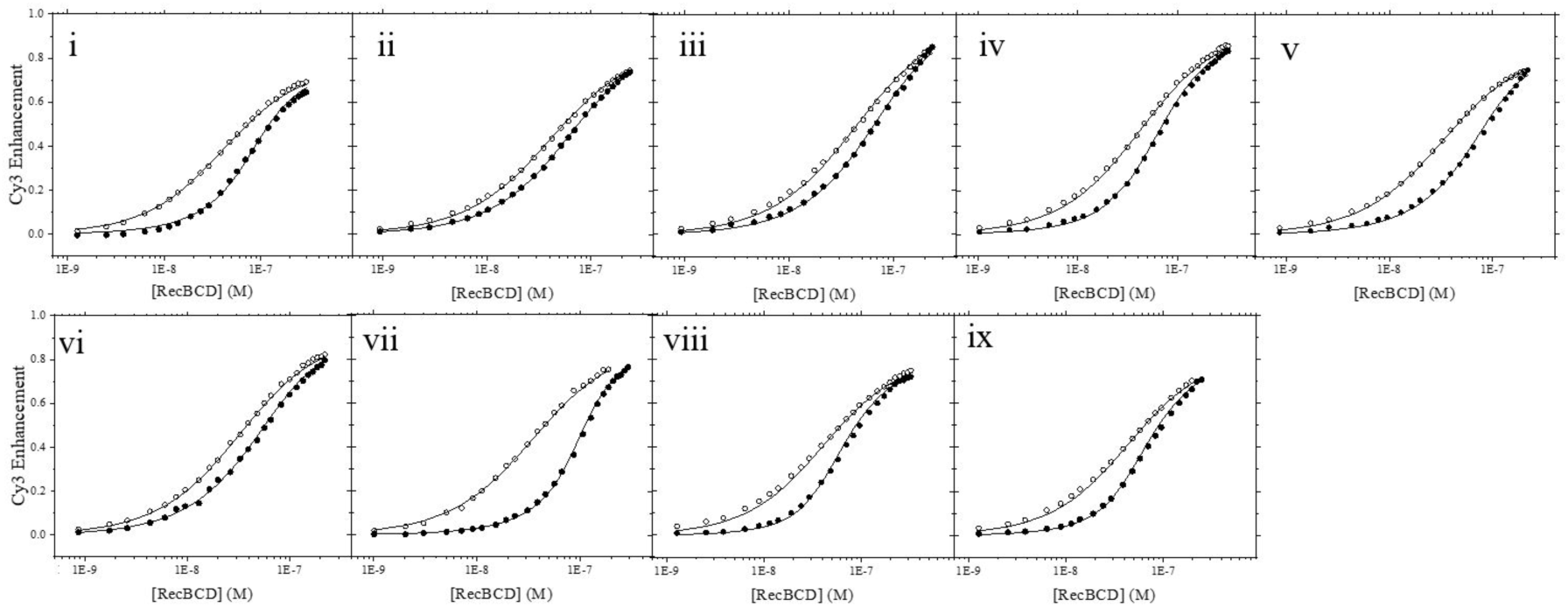
Binding isotherms from fluorescence competition titrations of RecBCD to either reference DNA (empty circle) or a mixture (solid circles) of Reference DNA and blunt (i), 3’-dT_2_ (ii), 3’- dT_4_ (iii), 3’-dT_6_ (iv), 3’-dT_10_ (v), 3’-dT_15_ (vi), 5’-dT_6_ (vii), 5’-dT_10_ (viii) or 5’-dT_15_ (ix). Solid lines indicates global fitting of two isotherms within each panel. Experiments were performed in Buffer M275-10, 25°C. Equilibrium constants summarized in Supplementary Table 2.

**Supplementary Figure 2.**
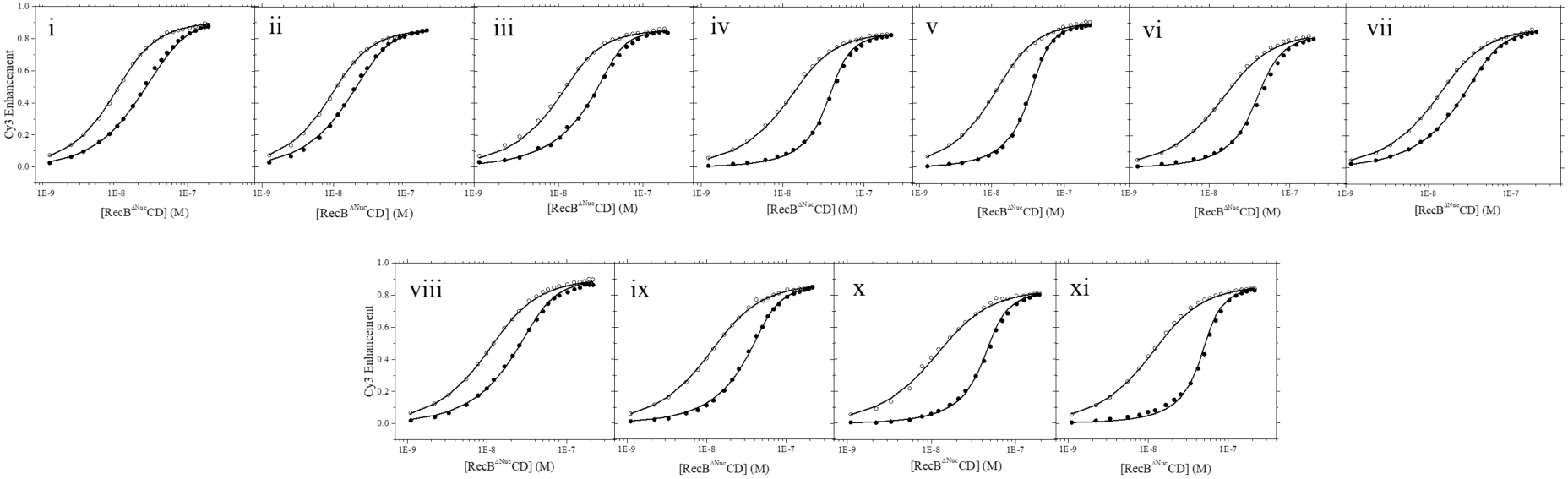
Binding isotherms from fluorescence competition titrations of RecB^ΔNuc^CD to either reference DNA (empty circle) or a mixture (solid circles) of Reference DNA and blunt (i), 3’dT_2_ (ii), 3’dT_4_ (iii), 3’dT_6_ (iv), 3’dT_8_ (v), 3’dT_10_ (vi), 3’dT_15_ (vii), 5’dT_6_ (viii), 5’dT_8_ (ix), 5’dT_10_ (x) or 5’dT_15_ (xi). Solid lines indicates global fitting of two isotherms within each panel. Experiments were performed in Buffer M275- 10, 25°C. Equilibrium constants summarized in Supplementary Table 2.

**Supplementary Figure 3.**
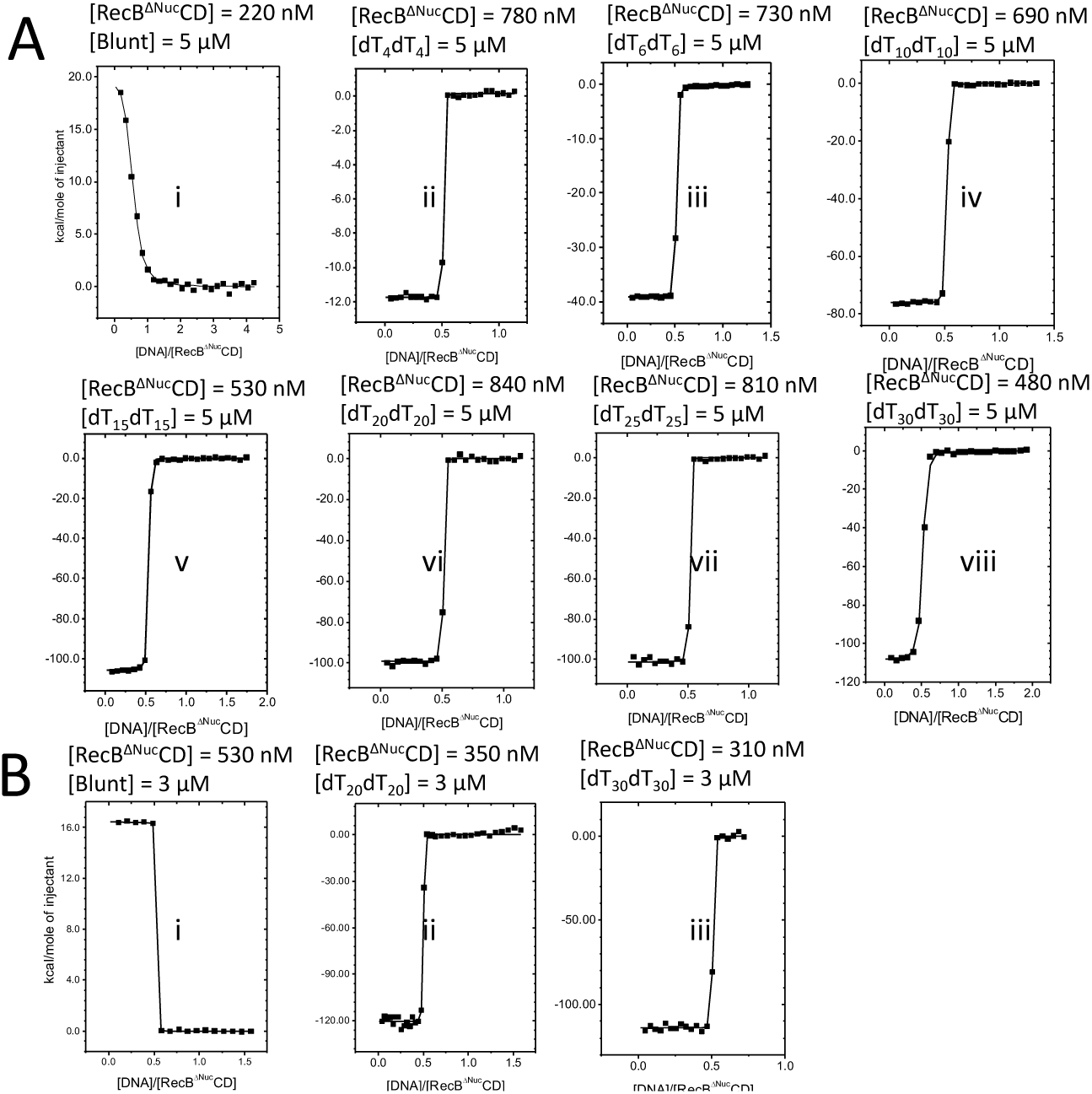
ITC binding isotherms for results in Figure 2D Concentrations of RecB^ΔNuc^CD and titrated DNA substrate are indicated above each panel for ITC experiments performed in Buffer M275-10, 25°C (A) or in cryo-EM buffer, 25°C (B). The thermodynamic parameters determined from these experiments are summarized in Supplementary Table 3. RecB CD binding to Blunt DNA in Buffer M275-10, 25°C has a binding constant of K=1.2(±0.4)×10^7^M^-1^. All the rest of the binding isotherms have binding constant that are too high to be accurately measured, with a lower limit of K>10^9^M^-1^.

**Supplementary Figure 4.**
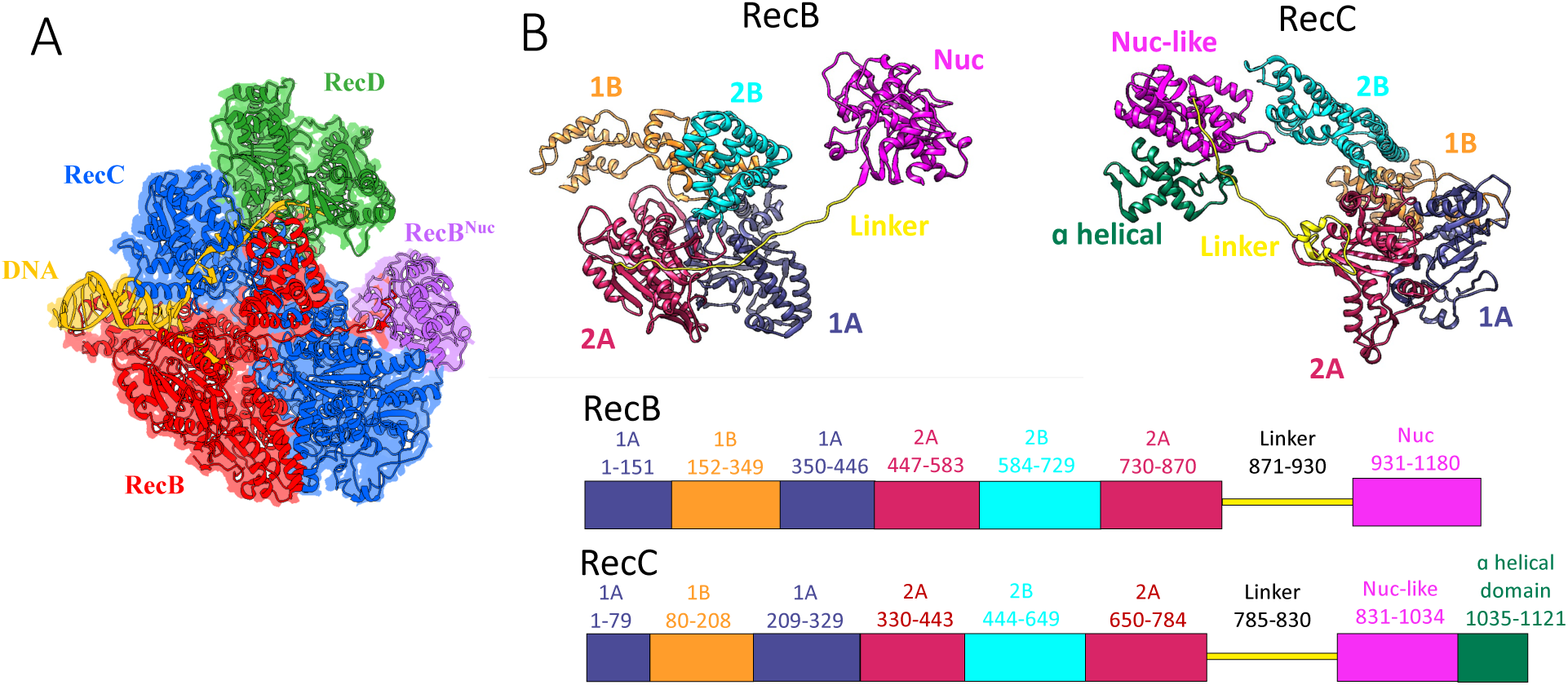
Supplementary Figure 1. (A) Cartoon depiction of RecBCD to aid visualization of relative positions of various components of the complex, with RecB motor domains in red, RecB^Nuc^ in purple, RecC in blue, RecD in green and DNA in yellow. (B) ribbon representations of RecB and RecC structures with various subdomains color coded with the linear representations below to aid visualization. The numbers on top of the linear representations indicate the amino acid numbers of each region.

**Supplementary Figure 5.**
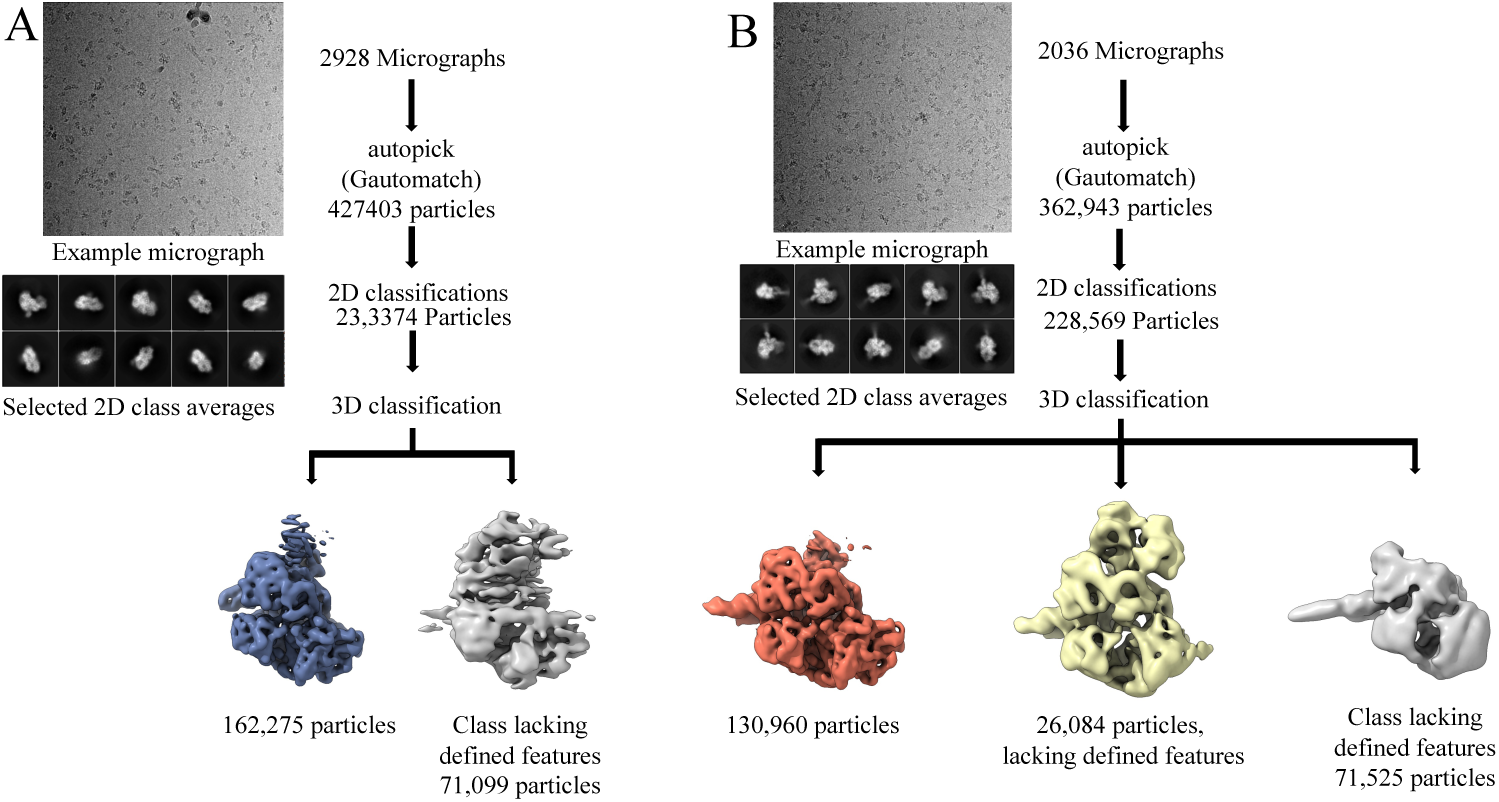
Workflows for single particle analysis of RecB^ΔNuc^CD (A) or RecB^ΔNuc^CD-DNA complex (B).

**Supplementary Figure 6.**
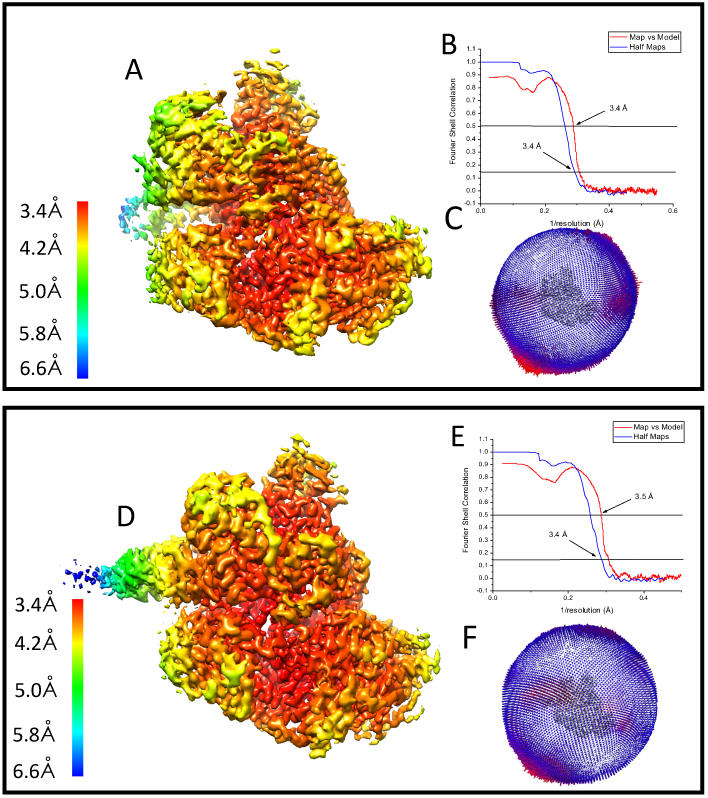
Cryo-EM data statistics **(A-C)** are statistics for RecB^ΔNuc^CD. **(D-F)** are statistics for RecB^ΔNuc^CD-DNA. **(A)** and **(D)** are estimates of local resolutions; **(B)** and **(E)** are overall resolutions; **(C)** and **(F)** are angular distributions of particles.

**Supplementary Figure 7.**
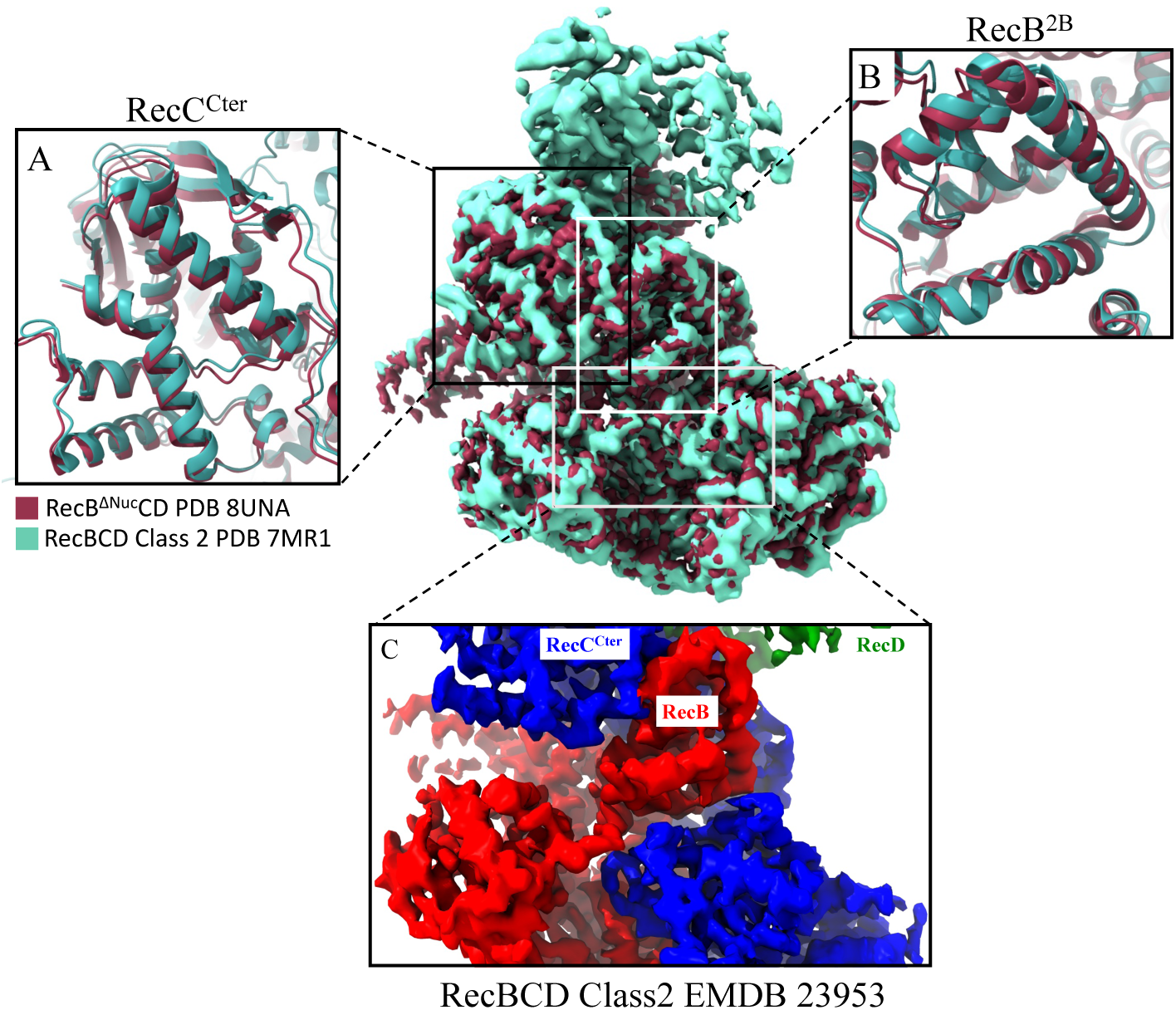
Cryo-EM maps of RecB^ΔNuc^CD (dark red, EMDB 42396) and RecBCD Class 2 (grey, EMDB 23953) were overlaid, and several regions of interests were compared. Panel A and B compare conformations of RecC^Cter^ and RecB^2B^ between RecB^ΔNuc^CD and RecBCD respectively. Panel C uses the cryo-EM map of Class 2 RecBCD as an example to show that the map densities representing RecB^Linker^ and RecC^Linker^ were not observed. This is the same for RecB^ΔNuc^CD.

**Supplementary Figure 8.**
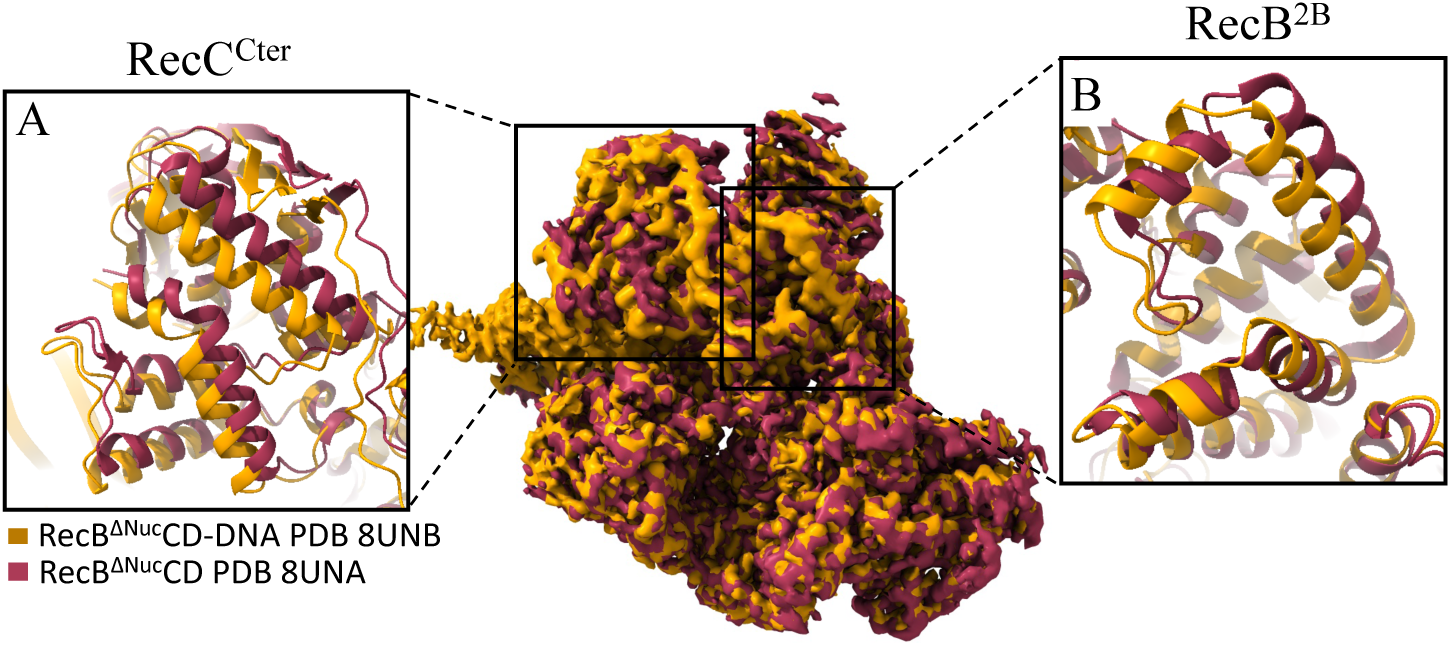
Structural comparison between RecB^ΔNuc^CD and RecB^ΔNuc^CD-DNA, demonstrating conformation shifts in RecC^Cter^ (A) and RecB^2B^ (B), despite RecB^Linker^ and RecC^Linker^ densities are absent in both structural maps.

**Supplementary Figure 9.**
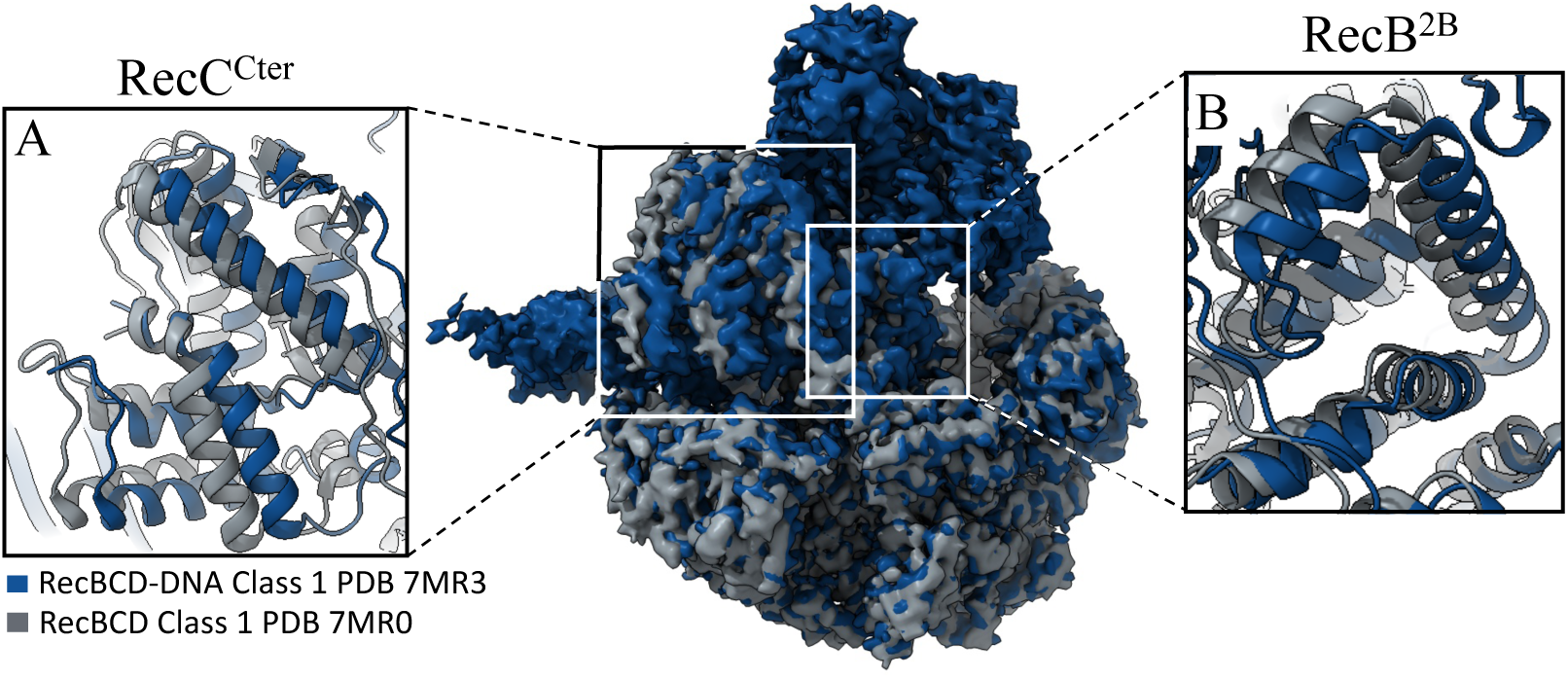
Structural comparison between RecBCD Class 1 and RecBCD-DNA Class 1, demonstrating conformation shifts in RecC^Cter^ (A) and RecB^2B^ (B).

**Supplementary Figure 10.**
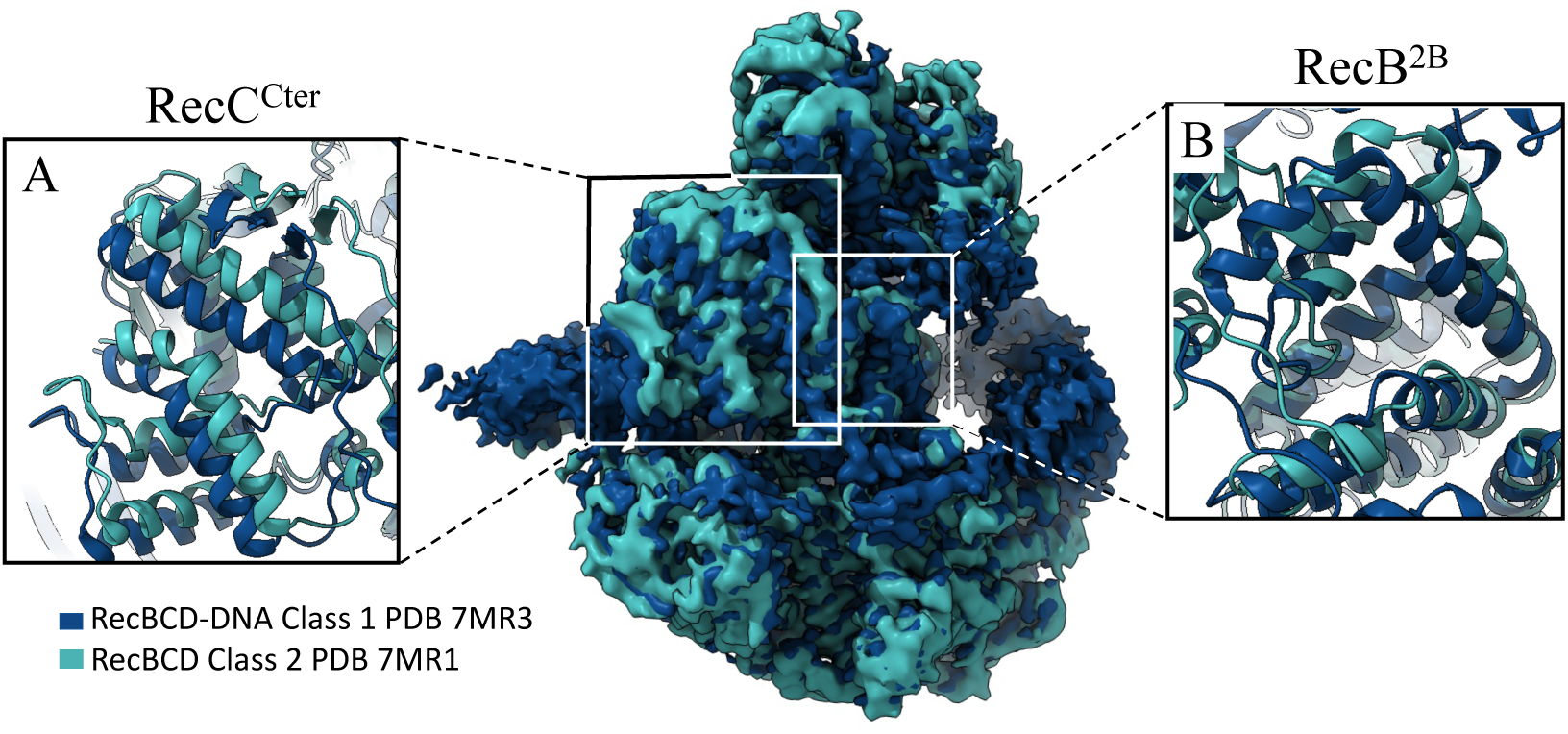
Structural comparison between RecBCD Class 2 and RecBCD-DNA Class 1, demonstrating conformation shifts in RecC^Cter^ (A) and RecB^2B^ (B).

